# Human HELB is a processive motor protein which catalyses RPA clearance from single-stranded DNA

**DOI:** 10.1101/2021.05.27.445972

**Authors:** S Hormeno, OJ Wilkinson, C Aicart-Ramos, S Kuppa, E Antony, MS Dillingham, F Moreno-Herrero

## Abstract

Human HELB is a poorly-characterised helicase suggested to play both positive and negative regulatory roles in DNA replication and recombination. In this work, we used bulk and single molecule approaches to characterise the biochemical activities of HELB protein with a particular focus on its interactions with RPA and RPA-ssDNA filaments. HELB is a monomeric protein which binds tightly to ssDNA with a site size of ∼20 nucleotides. It couples ATP hydrolysis to translocation along ssDNA in the 5′-to-3′ direction accompanied by the formation of DNA loops and with an efficiency of 1 ATP per base. HELB also displays classical helicase activity but this is very weak in the absence of an assisting force. HELB binds specifically to human RPA which enhances its ATPase and ssDNA translocase activities but inhibits DNA unwinding. Direct observation of HELB on RPA nucleoprotein filaments shows that translocating HELB concomitantly clears RPA from single-stranded DNA.

## INTRODUCTION

The human HELB protein, also known as human DNA helicase B (HDHB), was first identified as a homologue of a putative murine replicative helicase (Hazeslip et al., 2020; Saitoh et al., 1995; Taneja et al., 2002). Since then, various functions have been assigned to the protein including a role in the onset of chromosomal DNA replication (Taneja et al., 2002), cellular recovery from replication stress (Guler et al., 2012), promotion of Cdc45 chromatin-binding (Gerhardt et al., 2015), resolution of DNA secondary CGG repeat structures (Guler et al., 2018) and stimulation of RAD51-mediated 5′-3′ heteroduplex extension to promote homologous recombination (HR) (Liu et al., 2015). Most recently, and in apparent contradiction to the role in the stimulation of HR, HELB was proposed to inhibit homology-dependent double-stranded DNA break repair (DSBR) by antagonising the processive resection nucleases EXO1 and DNA2/BLM during the G0/G1 phases of the cell cycle (Tkáč et al., 2016). In agreement with this idea, HELB forms nuclear foci in response to DNA damage and is phosphorylated by cyclin-dependent kinase (CDK) causing localisation to the nucleus in G1 and to the cytoplasm during S/G2. The formation of HELB damage foci is dependent on the main eukaryotic single-stranded DNA binding protein Replication Protein A (RPA) (Wold, 1997), which has been shown to interact physically with HELB (Guler et al., 2012; Tkáč et al., 2016). Although the interaction with RPA is potentially critical for all putative functions of HELB, the ability of this motor protein to modulate the formation, remodelling or removal of RPA nucleoprotein filaments has never been studied and is the focus of the work presented here.

The filaments formed between RPA and ssDNA are critical intermediates in DNA replication, recombination and repair (Dueva and Iliakis, 2020; Maréchal and Zou, 2015; Zou et al., 2006). RPA not only shields ssDNA from nucleolytic degradation, it is also involved in the recruitment or exclusion of other factors from ssDNA, the regulation of DNA replication and repair, and the initiation of cell signalling cues that link these pathways to the cell cycle and its progression through checkpoints (Dhingra et al., 2021). Interestingly, many helicases and helicase-like proteins share intimate physical and functional interactions with single-stranded DNA binding proteins (Awate and Brosh Jr, 2017; Caldwell and Spies, 2020). However, we do not currently understand how the activity of HELB affects RPA filaments and *vice versa*.

HELB is a 120 kDa protein comprising three distinct domains: an N-terminal region of unknown function, a central helicase domain sharing homology with the Superfamily 1 (SF1) helicase RecD, and a C-terminal region containing CDK phosphorylation sites (Hazeslip et al., 2020) (**Figure 1A**). Site-directed mutagenesis has implicated the central helicase domain in both the DNA and RPA binding activities of HELB (**Figure 1A**, blue arrows). Interestingly, mutations in HELB are associated with both female infertility and early-onset menopause and are found widely distributed in the protein sequence in human tumour samples (**Figure 1A**, red arrows) (Day et al., 2015; Tate et al., 2019). *In vitro* studies show that HELB possesses single-stranded DNA-dependent ATPase activity and 5′-to-3′ helicase activity, which is as expected based on the similarity to RecD (Saikrishnan et al., 2008; Taneja et al., 2002). However, precisely how these biochemical properties underpin the cellular function(s) of HELB and the significance of the interaction with RPA are unresolved.

**Figure 1.**
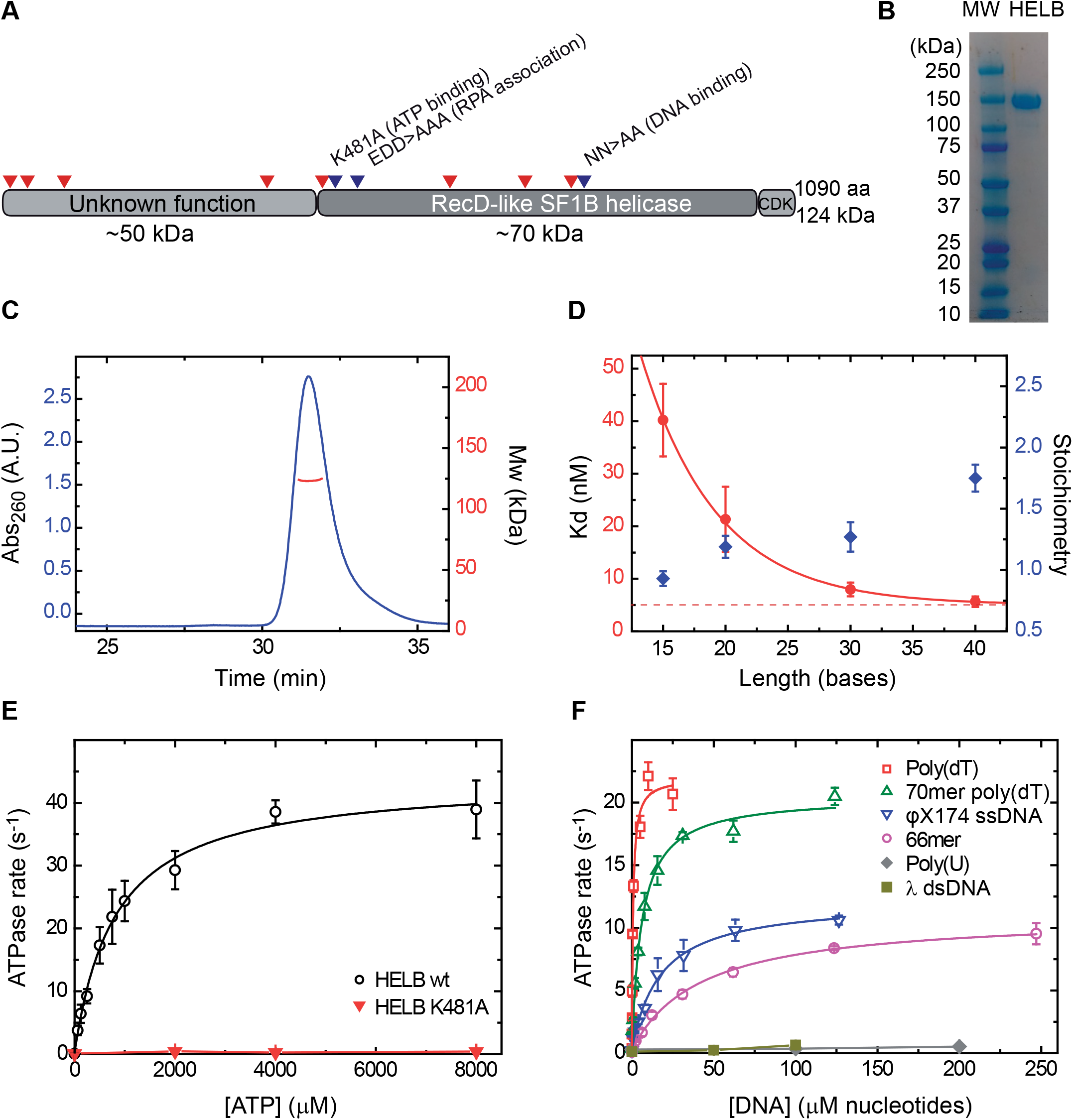
HELB is a monomer which binds tightly to single-stranded DNA and displays ssDNA-dependent ATPase activity. (**A**) Cartoon of HELB showing overall domain layout and important mutations. Red marks denote positions of high frequency tumour mutations. Blue marks denote positions of mutations that affect ATPase, DNA binding and RPA binding activities. (**B**) SDS-PAGE analysis shows highly-purified recombinant human HELB produced from insect cells. (**C**) SEC-MALS analysis demonstrates that HELB is a monomer in solution under these conditions with a calculated molecular weight of 123,252 Da. (**D**) (left axis), HELB binding constants (Kd) for Poly(dT) substrates of different lengths obtained in PIFE assays described in **Figure S1A**. An exponential fit determines a saturating Kd of 5 nM. (Right axis), stoichiometry values obtained under tight binding conditions as shown in **Figure S1B-E**. (**E**) Michaelis-Menten plot of ATP hydrolysis gives *K*_m_ and *k*_cat_ parameters for WT HELB, and also shows that the K481A mutant is unable to hydrolyse ATP. (**F**) Analysis of DNA stimulation of HELB ATPase activity demonstrates that HELB is an ssDNA-dependent helicase. ATP turnover is stimulated more by polythymidine substrates than mixed-base sequences, likely due to their inability to form inhibitory secondary structures.

In this study, we used bulk and single-molecule assays to further characterise HELB, including its physical and functional interactions with RPA and RPA nucleoprotein filaments. Paradoxically, we find that RPA (itself a potent ssDNA binding protein) is stimulatory to all activities of HELB on ssDNA despite the competition one would expect between the two proteins for their nucleic acid substrates. In contrast, non-cognate single-stranded DNA binding proteins inhibit all activities of human HELB. These highly specific interactions with RPA filaments help to recruit HELB onto ssDNA that is devoid of secondary structure and promote efficient ssDNA translocation coupled to the processive clearance of RPA. The implications of this finding for the roles of HELB in DNA replication and recombination are discussed.

## RESULTS

### HELB is a monomeric protein which binds tightly to single-stranded DNA and displays ssDNA-dependent ATPase activity

Human HELB contains a C-terminal Superfamily I helicase domain and a large N-terminal region of unknown function which displays no apparent homology to known proteins or folds (**Figure 1A**). In order to better characterise this protein, we first prepared pure recombinant HELB from insect cells (**Figure 1B**). SEC-MALS analysis showed that native HELB has a molecular weight of 123 ± 3 kDa which is the expected value for a monomer (**Figure 1C**). To quantitatively investigate ssDNA binding activity, we used a protein-induced fluorescence enhancement (PIFE) assay to measure binding to a series of poly(dT) oligonucleotides of increasing length (**Figure S1A**). The affinity of HELB for ssDNA increased with increasing length until saturation at K_d_ ∼5 nM for oligonucleotides of 30 bases or longer (**Figure 1D**), suggesting a binding site size of between 20 and 30 nucleotides. We then performed the PIFE assay under tight binding conditions (i.e. using a DNA concentration above the measured K_d_) so as to calculate the apparent binding stoichiometry. We find that one HELB monomer binds to ssDNA molecules as long as 30 bases, whereas two HELBs can be accommodated by a 40mer oligonucleotide **(Figure 1D** and **S1B-E)**. Taken together, these data suggest a binding site size of approximately 20 nucleotides. This is much larger than is typical for a SF1 DNA helicases; the core helicase domains bind ∼8 nucleotides such that two monomers can bind side-by-side to a 16mer oligonucleotide (Gilhooly et al., 2013; Korolev et al., 1997). This in turn implies that HELB contains an additional undefined DNA binding site, perhaps in the N-terminal region of the protein, an idea which is supported further by experiments presented below.

HELB displays ssDNA-dependent ATPase activity with Michaelis-Menten parameters of <u>k</u>_cat_ = 44 ± 3 s^-1^ and *K*_m (ATP)_ = 800 ± 140 µM measured in the presence of saturating poly(dT) concentrations (**Figure 1E**). This turnover number is significantly higher than has been reported previously (280 ATP min^-1^) (Taneja et al., 2002). As expected, substitution of the conserved lysine in the Walker A motif (helicase motif I) to alanine (K481A) dramatically reduced ATPase activity showing that this activity is intrinsic to the purified HELB polypeptide (**Figure 1E**). We compared the ability of six model nucleic acid substrates to stimulate the ATPase activity of HELB (**Figure 1F**). Duplex DNA and poly(U) single-stranded RNA did not stimulate the ATPase rate. In contrast, single-stranded DNA strongly activated ATPase activity. We found that poly(dT) (a mixture of polythymidine chains of average length ∼1000 nucleotides) was a significantly better substrate for HELB than φX174 virion ssDNA (a circular 5386 nucleotide molecule), reflected both in terms of a higher *k*_cat_ and a lower *K*_DNA_. This indicates that secondary structure and/or DNA sequence may affect both the binding of HELB onto DNA and the DNA-stimulated ATPase that is coupled to translocation.

### HELB translocates efficiently on ssDNA in a 5′-to-3′ direction

To assess the putative ssDNA translocase activity of HELB we first used an indirect gel-based assay based on the displacement of streptavidin from biotinylated oligonucleotides (Morris and Raney, 1999). By comparing streptavidin displacement from oligonucleotides labelled at either the 3′-or the 5′-end with biotin, one can infer DNA translocation activity as well as its polarity. We observed that HELB efficiently removed streptavidin from 3′-biotin (but not 5′-biotin) labelled substrates, suggesting that HELB moves in the 5′-to-3′ direction **(Figures 2A** and **2B)**. This is the expected polarity given the sequence similarity to the RecD-family of helicases.

**Figure 2.**
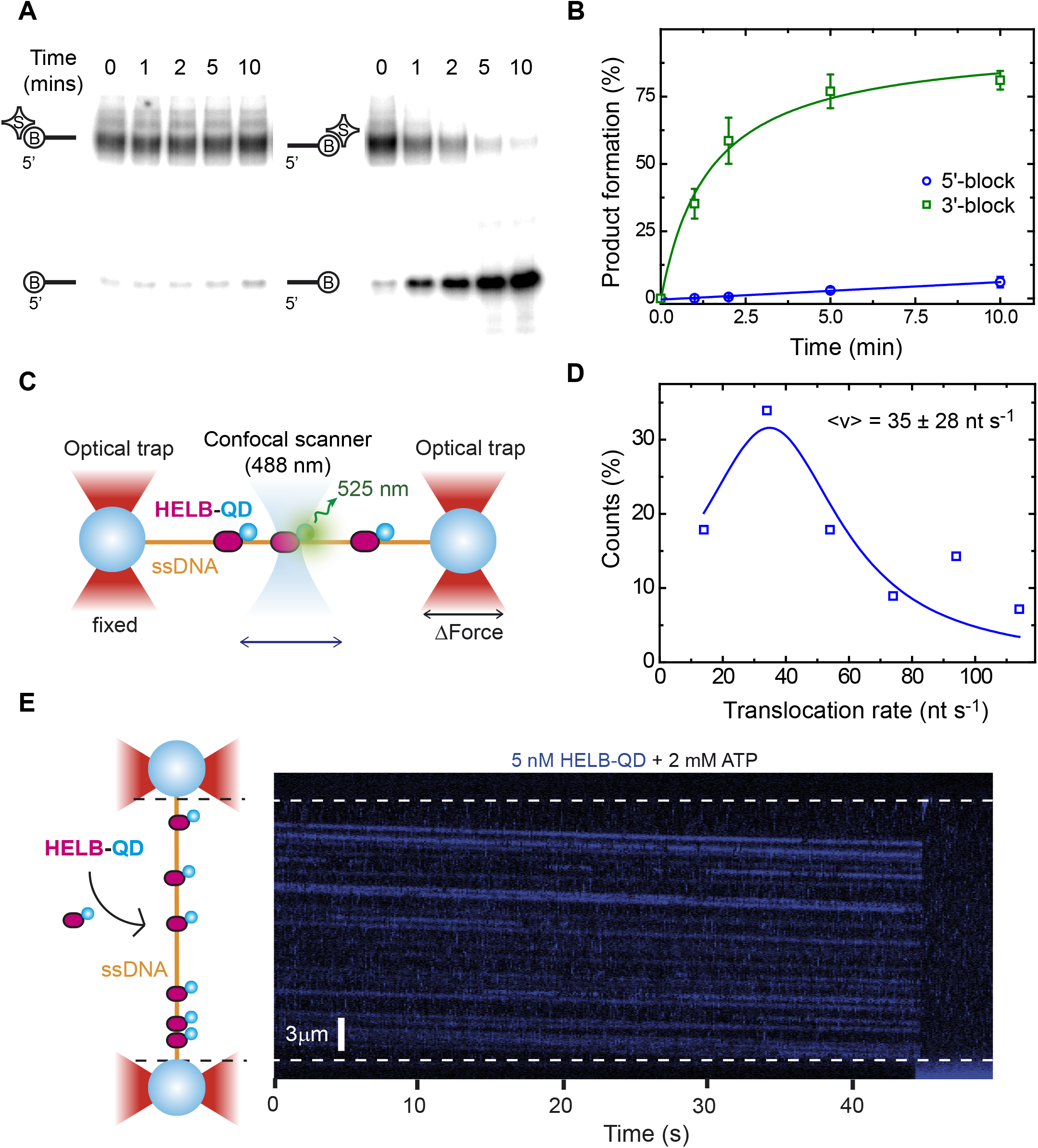
HELB efficiently translocates on ssDNA in a 5′-to-3′ direction. (**A**) Streptavidin displacement assay in bulk demonstrates that HELB moves along ssDNA specifically in a 5′-to-3′ direction. (**B**) Quantification of the gel-based assay. (**C**) Illustration of the experimental C-trap setup. Individual λ-sized ssDNA tethers were attached between two streptavidin-coated beads trapped by two optical traps. A confocal laser scanned the DNA tethers to detect HELB conjugated to one QD. (**D**) Distribution of HELB translocation rates on ssDNA hold at 14 pN in the presence of 2 mM ATP (n = 50). A Lorentzian fit has been included to guide the eye. (**E**) Cartoon of a tethered ssDNA loaded with multiples HELB-QDs (left) and representative kymograph of HELB movement (blue) in the presence of 2 mM ATP under 19 pN of tension (right).

Next, to better characterise the rate and processivity of ssDNA translocation, we directly imaged the movement of fluorescent HELB on single-stranded DNA using a combined optical tweezers and confocal fluorescence microscope (Candelli et al., 2011; Newton et al., 2019) (C-Trap, Lumicks). For these experiments, we first conjugated biotinylated HELB (which retains wild type ATPase activity; data not shown) to streptavidin-coated quantum dots (QDs). A long ssDNA substrate was generated from a λ phage dsDNA (48.5 kbp) that had been tethered between two optically trapped beads by applying a tension above the overstretching force in a low salt buffer (**Figure 2C** and **Figure S2A-B**). Successful generation of a ssDNA molecule was assessed by its mechanical fingerprint, which is very different from dsDNA (**Figure S2C**). Once a single ssDNA molecule was stably trapped between the beads, we moved the molecule into a channel containing 5 nM HELB-QD and 2 mM (saturating) ATP and recorded confocal images between the beads at 50-100 ms line^-1^, allowing us to build kymographs depicting HELB binding and movement on DNA (**Figure 2E**). These experiments showed that HELB binds to ssDNA and then translocates in a unidirectional fashion (**Figure 2E**). Because the ssDNA tether orientation is arbitrary, we cannot determine the translocation polarity and found examples of kymographs with HELB moving towards either bead (**Figure 2E and Figure S2D**). However, based upon our bulk experiments, movement is presumably in the 5′-to-3′ direction. In the absence of ATP, HELB was able to bind to ssDNA but remained stationary (**Figure S2E**). The rate of translocation on naked ssDNA at room temperature was 35 ± 28 nt s^-1^ (peak ± width/2, n = 56) (**Figure 2D**). This value is similar to the ATPase rate under similar conditions suggesting that HELB hydrolyses ∼1 ATP molecule per base translocated in common with other SF1 helicases (Gilhooly et al., 2013). Together, these experiments show that HELB is a processive motor protein which couples ATP hydrolysis to 5′-to-3′ unidirectional translocation along ssDNA without the need to initiate from a free DNA end.

### HELB is an efficient helicase only when assisted by force

To assess the extent to which HELB couples its ssDNA translocation activity to helicase activity (i.e., strand unwinding and separation), we first performed bulk helicase assays using a short duplex DNA containing a 5′-overhang as a loading site (**Figure 3A**). At elevated protein concentrations (500 nM) wild type, but not ATPase-dead mutant HELB, partially unwound the duplex revealing intrinsic helicase activity as reported previously (Taneja et al., 2002). However, the apparently weak helicase activity detected on this substrate contrasts with the potent translocase activity reported above. We next performed magnetic tweezers (MT) experiments to investigate further the kinetics of duplex unwinding by HELB and the potential role of the force in this activity. We fabricated a ∼6.3 kbp DNA substrate containing a 5′-terminated poly(dT) ssDNA (37 nt) positioned 1.7 kbp from one end that acts as a loading site for HELB, named as Flap-DNA (**Figure 3B**). One DNA end was attached to a glass surface of a fluid cell, and the other to a superparamagnetic bead. External magnets located above the cell were then used to apply a controlled force to extend the DNA while the height of the bead was monitored (**Figure 3C**). In this set-up, it is possible to monitor DNA unwinding because single-and double-stranded DNA display different extension for a given force (Dessinges et al., 2004). In the high applied-force regime (F ≥ 6 pN), ssDNA is longer than duplex DNA and so helicase activity leads to an increase in the Z position of the bead. Under low forces (F < 6 pN), single-stranded DNA is shorter than duplex and unwinding leads to a reduction in the height of the bead (Bustamante et al., 2003; Carrasco et al., 2020).

**Figure 3.**
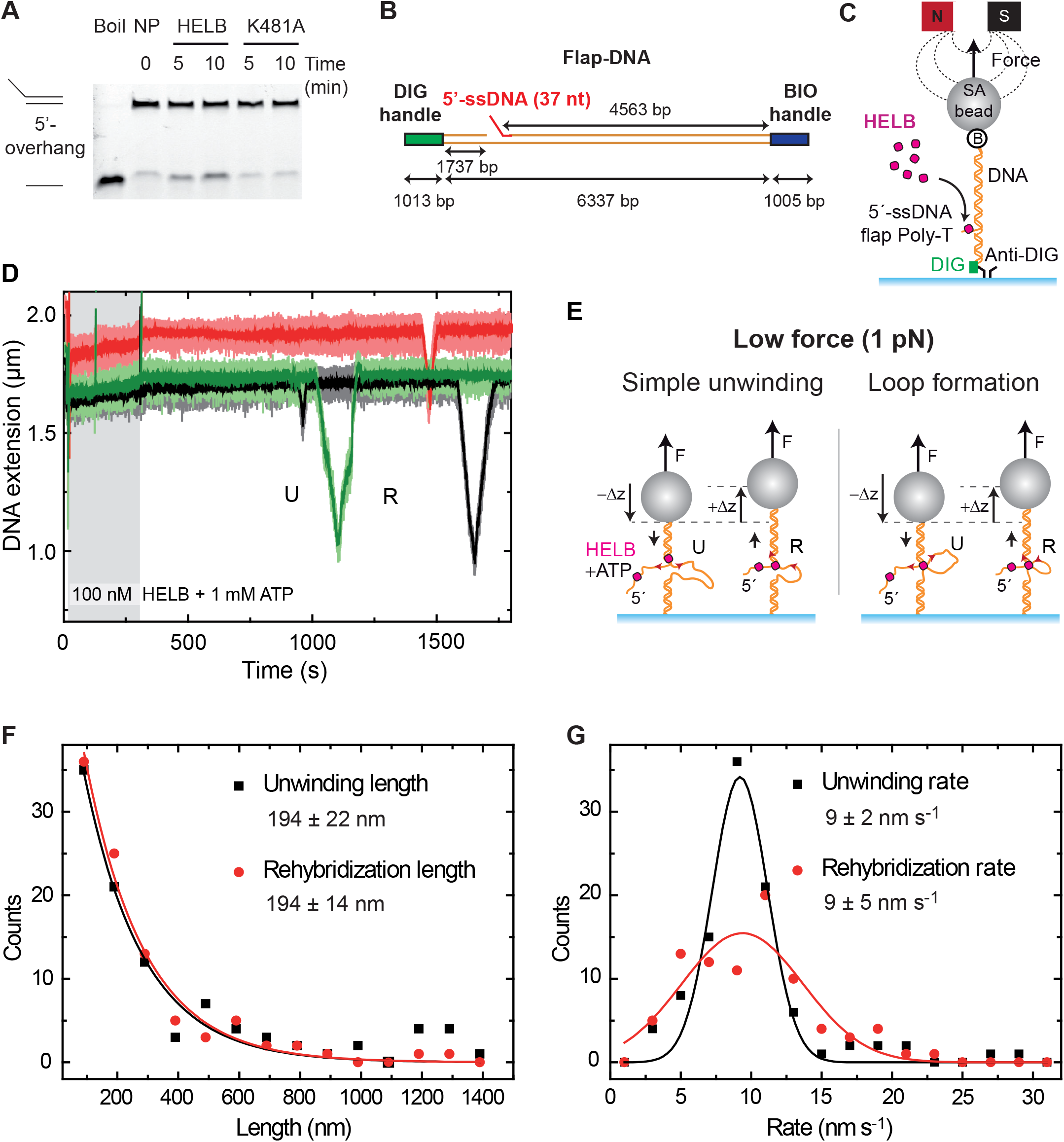
HELB can unwind duplex DNA and switch translocating strands. (**A**) Bulk unwinding assay shows limited HELB duplex-unwinding activity. (**B**) Schematic representation of the Flap-DNA substrate employed for MT single-molecule unwinding assays. (**C**) Cartoon of the MT assay for monitoring DNA unwinding by HELB. (**D**) Representative time-courses of MT experiments performed at 1 pN with 100 nM HELB and 1 mM ATP. Several events where the DNA extension decreases (U) and then increases again (R) to reach the initial bead’s height are shown. (**E**) The U events can be explained by direct HELB unwinding at low forces (exposing a stretch of ssDNA) or by loop formation. Upon strand switching, the HELB translocation along the opposite strand leads to the rehybridization of the helix and the recovery of the initial height of the bead (R events). (**F**) Distribution of the unwinding length (black) and rehybridization length (red) measured in MT time-courses. Fittings to two exponential decays give the mean values of <L_U_> = 194 ± 22 nm (error of fitting, n = 99) and <L_R_> = 194 ± 14 nm (error of fitting, n = 94). (**G**) Distribution of the unwinding rate (black) and rehybridization rate (red) of events extracted from MT time-courses similar to those shown in **D**. Fits to Gaussian functions give the mean unwinding rates of <v_U_> = 9 ± 2 nm s^-1^ (n = 99) and <v_R_> = 9 ± 5 nm s^-1^ (n = 89).

MT unwinding experiments were first performed in the low applied-force regime (1 pN). Upon addition of 100 nM HELB and saturating ATP, we observed cycles of unwinding (U), reflected in a reduction of the bead height, and rehybridization (R) events, in which the initial extension was recovered (**Figure 3D**). A control gap-substrate was also prepared containing a 63-nt-gap and no flap, named as Gap-DNA (**Figure S2F**). No activity was observed in control experiments using the Gap-DNA substrate, suggesting that unwinding initiates from the free 5′-ssDNA tail (**Figure S2G**). To estimate the number of unwound base pairs and the velocity, we measured the change in extension in time and considered two models (**Figure 3E**). The first is based on the formation of expanding ssDNA regions (i.e. simple unwinding). The second, which requires an additional DNA binding site and is included for reasons that will become more apparent below, is based on ssDNA loop formation. In the simple unwinding scenario, HELB would have unwound 1240 ± 140 bp (error of the exponential fit, n = 99) at an unwinding rate of 56 ± 11 bp s^-1^ (peak ± w/2, n =99). In the ssDNA looping model, HELB would have unwound 865 ± 146 bp at 45 ± 10 bp s^-1^; a rate very similar to that observed in the ATPase and ssDNA translocase assays above. Regardless of the model, the rehybridization parameters were similar to those obtained in unwinding time-courses (**Figure 3F** and **3G**). During unwinding, we usually observed a uniform decrease of the DNA extension without pauses, while rehybridization occurred punctuated by pauses or even backtracking. On the other hand, the similar unwinding and rehybridization rates, as well as the apparent symmetry of these events, support the idea that they reflect strand switching by a single HELB enzyme (Dessinges et al., 2004). We conclude that HELB can processively unwind DNA at low force. It is important to note, however, that these events were extremely infrequent, as they were typically observed several minutes after injection of HELB (**Figure S3A**). These rare events observed at the single molecule level at low force are consistent with bulk DNA helicase assays (at zero force) which detect only a very limited 5′-to-3′ helicase activity (**Figure 3A**). Further control experiments performed in the absence of ATP (**Figure S3B**), with the ATPase-dead mutant K481A (**Figure S3C**), and on nicked (**Figure S3D**) or torsionally-constrained DNA substrates (not shown) did not show any helicase activity.

We next performed equivalent magnetic tweezers experiments but at a higher applied-force of 8.4 pN, conditions under which ssDNA is longer than duplex DNA (**Figure 4A**). We typically observed a rapid elongation of the tethers consistent with HELB efficiently unwinding DNA (representative time-courses are shown in **Figure 4B**). The time required to observe any activity (activation time) for these events was exponentially distributed with a time constant of 147 ± 10 s (error of fitting, n = 44), much shorter than that observed in the low force regime (**Figure S3E**). Moreover, this value corresponds to (and is therefore limited by) the arrival of the protein and ATP to the position of the tracked tethers in the middle of the fluid cell. Helicase activity was observed in 48/74 traces (65%) while 35% showed no change in tether extension which we attribute to the potential absence of the 5′-flap in the DNA substrates. Additional control experiments performed at high force did not show any helicase activity (**Figure S3F-H**). Under these high force conditions, some unwinding of the Gap-DNA substrate was observed but the activation time was increased ∼4-fold compared to the Flap-DNA substrate (not shown). Otherwise, the substrate displayed similar unwinding length and unwinding rate parameters (not shown). We interpret these activities as starting from the 5′-end at the gap when it is partially melted by the high force applied. The observed unwinding length at 8.4 pN was 3600 ± 900 bp (peak of the distribution ± width/2, n = 48) (**Figure 4C**), which implies that HELB can translocate and unwind the entire substrate. The unwinding rate distribution provided a mean value of 38 ± 4 bp s^-1^ (peak ± w/2, n = 47) (**Figure 4D**), which is very similar to both the ssDNA translocation rate obtained in C-Trap experiments (**Figure 2D**) and the ATPase rate measured in bulk (**Figure 1E**). Overall, we interpret these traces as showing that HELB efficiently couples ATP hydrolysis to helicase activity that is initiated from the flap in the 5′-to-3′ direction when a high assisting force has been applied. Improved DNA unwinding with increasing force has been predicted as a general feature of other helicases (Pincus et al., 2015).

**Figure 4.**
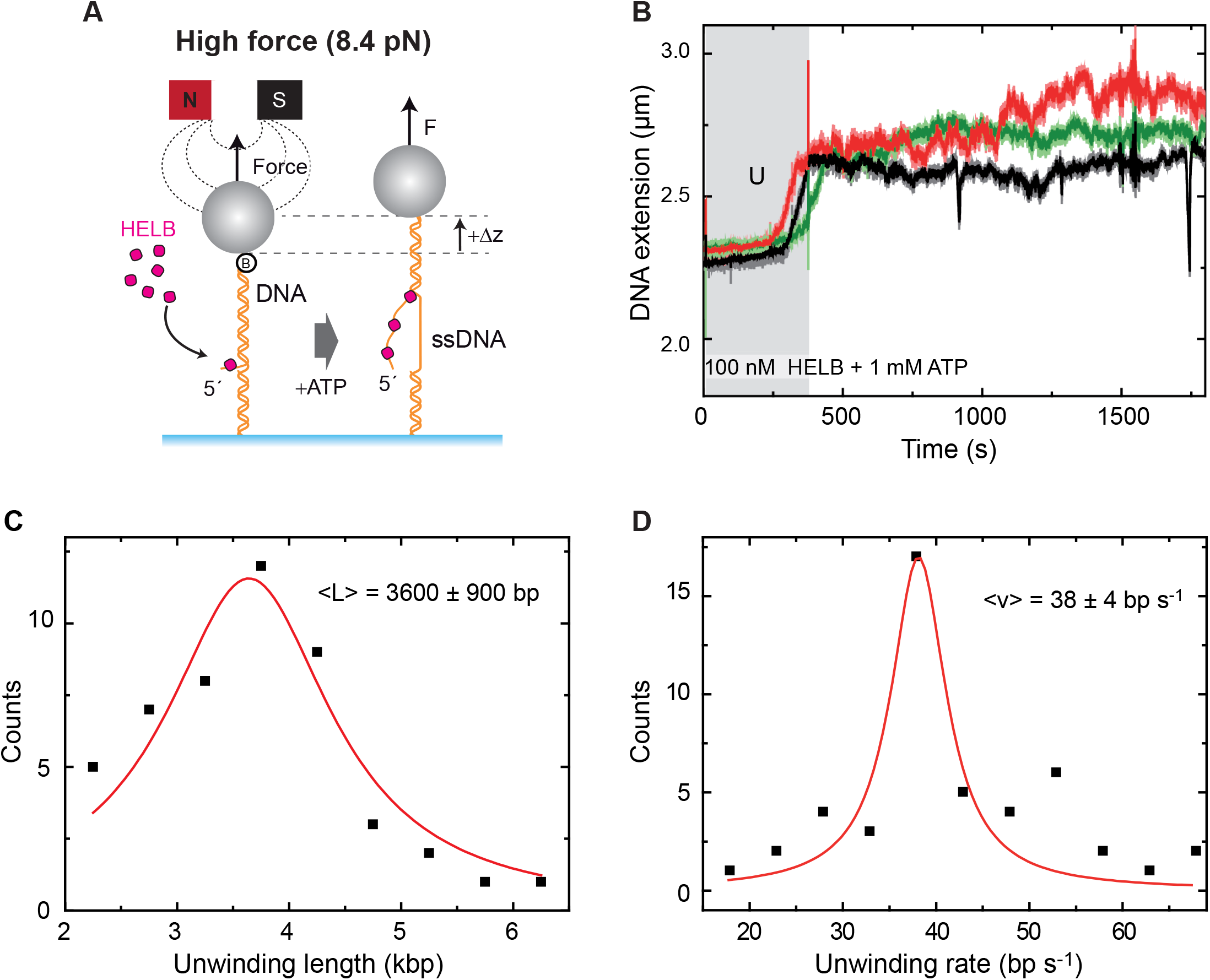
HELB efficiently unwinds duplex DNA when assisted by force. (**A**) MT model to explain duplex unwinding at high force (i.e., F ≥ 6 pN) using the Flap-DNA substrate. At this force the extension of ssDNA is larger than that of dsDNA, and HELB unwinding results in an increase in the height of the bead. (**B**) Representative time-courses of 100 nM HELB, 1 mM ATP taken at 8.4 pN, showing characteristic unwinding (U) events. (**C**) Distribution of the unwinding length of events fitted to a Lorentzian function (red) with a mean unwinding length of <L> = 3600 ± 900 bp (n = 48). (**D**) Distribution of the unwinding rate of events fitted to a Lorentzian function with a peak at 38 ± 4 bp s^-1^ (n = 47).

### HELB translocation can result in the formation of DNA loops

Following the rapid and processive DNA unwinding, our high force time-courses presented a more complex behaviour with continuous changes in extension with no obvious directionality (**Figure 4B**). Occasionally we noticed that, despite the high restraining force, the bead’s height dropped rapidly followed by a recovery of the extension (**Figure S4A**). Events of this type occurred in about half of our long time-courses. Because ssDNA is longer than duplex under these conditions (Dessinges et al., 2004), these events could be caused either by the conversion of ssDNA into dsDNA (i.e. annealing activity), or by the generation of a DNA loop (Wilkinson et al., 2020). The first option would imply a strand switch of the unwinding HELB enzyme followed by the reannealing of the previously-separated strands. However, this scenario can be discounted because we clearly observe the bead proceeding to below the original height of the tether (**Figure S4A**). We therefore favour the second explanation, whereby the activity of HELB leads to the formation of large DNA loops. Such behaviour requires that the enzyme oligomerizes and/or that the monomer has more than one DNA binding locus (**Figure S4B**). We observed two different kinds of height recovery following looping: a sudden jump in the bead’s height (in 18 of 40 events) (not shown) or a gradual recovery of its initial position (in 22 of 40 events), including some pausing in discrete steps (**Figure S4A**). Both behaviors are consistent with the looping model depending on the nature of translocating blockage that cause the generation of the loop and how that is resolved. The length of the unwinding events that result in the formation of a loop (looping length) measured at 8.4 pN was 438 ± 60 bp (error of the exponential fit, n = 40) (**Figure S4C**) and the looping rate was 33 ± 14 bp s^-1^ (peak ± w/2, n = 37) (**Figure S4D**), similar to the rate of unwinding measured in the same experimental conditions (**Figure 4D**).

### Interaction of HELB with RPA-coated DNA increases ATPase activity and favours ssDNA loop formation

A direct interaction between HELB and RPA has been reported previously (Guler et al., 2012). However, using blue native PAGE **(Figure 5A)** and SEC (not shown), we were only able to detect a complex between RPA and HELB in the presence of ssDNA. EMSA experiments using a 5′-Cy5-labelled 25mer oligonucleotide demonstrated that HELB was able to supershift the RPA:DNA complex (**Figure 5B**). Although this oligonucleotide is too short to fully accommodate both proteins side-by-side (RPA binds 20-30nt, HELB binds 20-25nt) we cannot exclude the idea that the apparent protein:protein interaction we see here is mediated by the DNA. Importantly however, the ternary complex formed was species-specific as no supershift was evident when substituting human RPA for *S. cerevisiae* RPA (yRPA) **(Figure 5B)**.

**Figure 5.**
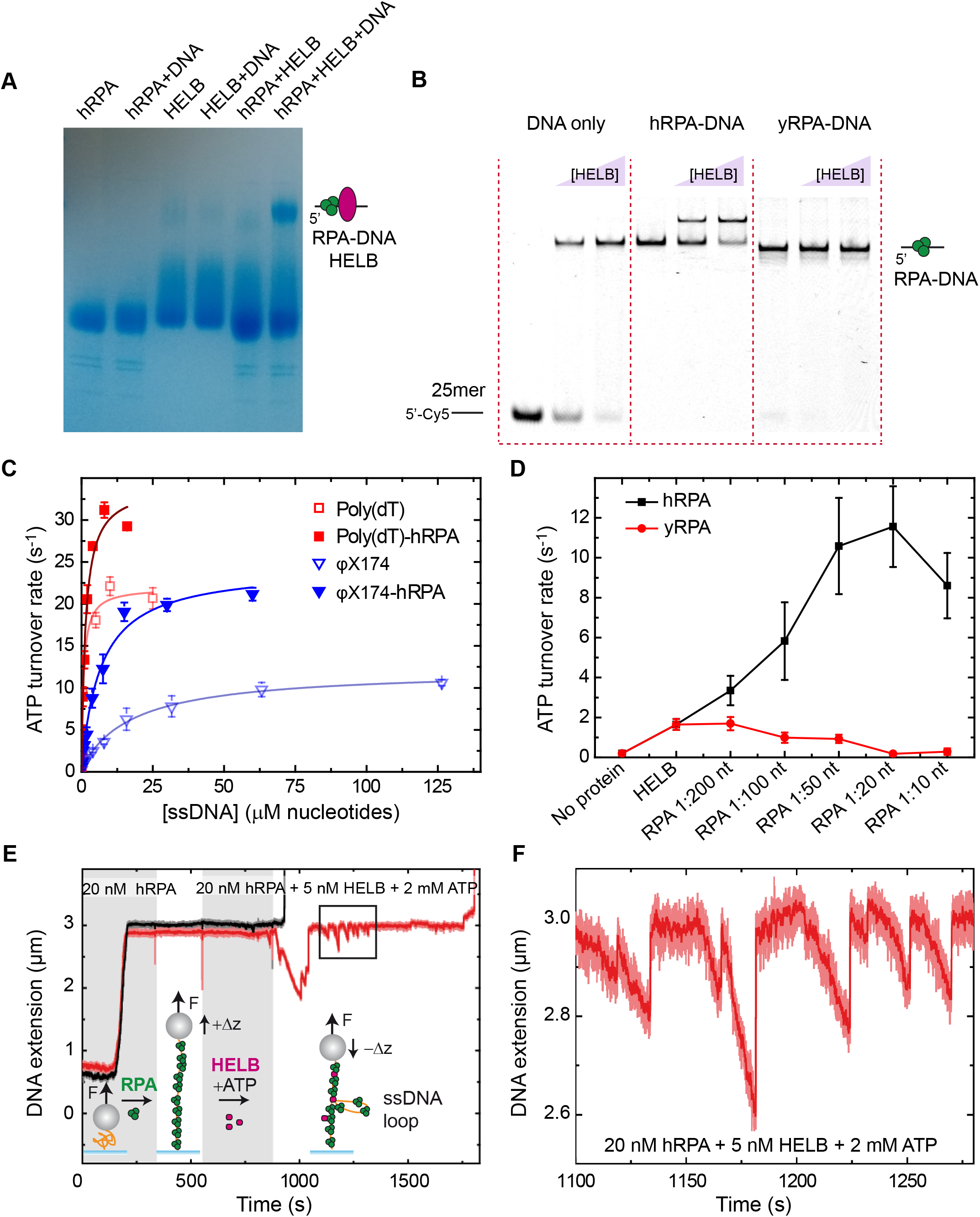
Interaction of HELB with RPA-coated DNA increases ATPase activity and favours ssDNA loop formation. (**A**) Native blue gel analysis confirms the formation of a HELB-human RPA-ssDNA complex with no apparent interaction observed for HELB and RPA without ssDNA. (**B**) Native–PAGE EMSA analysis of HELB interaction with ssDNA, human RPA-coated ssDNA, and yeast RPA-coated ssDNA shows specific super-shifted complex formation between HELB and human RPA-DNA. (**C**) Addition of RPA-ssDNA modifies HELB ATPase activity on the model substrates Poly(dT) and Virion φX174 (faded data is the same shown in **Figure 1F**). In both cases *k*_cat_ increases, but for the mixed base substrate, *K*_DNA_ also significantly decreases. (**D**) Stimulation of ATPase activity is unique to human RPA as yeast RPA causes inhibition of ATPase activity. (**E**) Example of a MT experiment where two ssDNA tethers covered with human RPA are measured under the effect of 5 nM HELB and 2 mM ATP (F = 3.8 pN). Shadow time-windows indicate the flow of RPA, and RPA+HELB+ATP. (**F**) Zoom-in of the square area marked in **E** to highlight the ssDNA looping dynamics promoted by HELB.

We next investigated the effect of RPA on the ATPase parameters of HELB using two model ssDNA substrates: poly(dT) which is comprised of long tracts of thymidine and incapable of forming secondary structures, and φX174 virion DNA which is expected to form extensive secondary structures **(Figure 5C)**. As shown above **(Figure 1F)**, in the absence of RPA, Virion φX174 is a much poorer substrate for HELB (lower *k*_cat_ and dramatically higher *K*_DNA_ compared to poly(dT)). We interpreted this as reflecting the high secondary structure content in virion DNA causing a physical block to binding and translocation. Naively, one might expect RPA (a tight ssDNA binding protein) to inhibit the ssDNA-dependent ATPase activity of HELB due to simple competition. However, we observed the opposite: substituting naked DNA for pre-bound DNA-RPA complexes (initially at 1 RPA:200 nucleotides, well-below saturation) resulted in an increase in *k*_cat_ for both substrates. This implies that the presence of RPA on DNA is somehow stimulating the ATPase (and possibly the translocation activity) of HELB in a manner unrelated to secondary structure content, and potentially due to the direct physical interaction. Strikingly, and selectively for the virion DNA, addition of RPA also resulted in a marked decrease in *K*_DNA_ (i.e. apparently tighter binding). The fact that this did not occur with poly(dT) implies that RPA is facilitating recruitment of HELB to DNA by resolving DNA secondary structures that are otherwise unfavourable for binding.

To investigate whether stimulation of HELB ATPase activity by RPA was dose-dependent we performed an RPA titration experiment at a low concentration of X174 ssDNA (0.1 × *K*_DNA_) where the observed ATPase rate is highly sensitive to stimulation **(Figure 5D)**. As RPA concentration increases from 1 RPA per 200 nt to 1 RPA per 20 nt (at which point we expect RPA to saturate the ssDNA), the observed rate of ATP hydrolysis increases by almost 8-fold. At concentrations above saturation (1 per 10 nt) there is a modest decrease from the maximum, suggesting that free RPA in solution is inhibitory to HELB ATP turnover. In complete contrast to the situation with human RPA, equivalent titrations with yeast RPA strongly inhibit HELB ATP turnover, presumably due to a simple competition for their substrates. This dose-dependent effect of RPA concentration and its species-selectivity on HELB activity was also observed in bulk translocation assays (**Figure S5A-B**). Taken together, these experiments show that the cognate RPA specifically recruits and stimulates the ATPase activity of HELB on ssDNA, implying that RPA nucleoprotein filaments are the physiological substrate for HELB.

Next, we investigated the interaction of HELB with RPA filaments using single molecule techniques. We first prepared ssDNA tethers for MT experiments following the methodology described in (Carrasco et al., 2020). Briefly, two strands of a dsDNA substrate are heat denatured followed by rapid cooling to avoid rehybridization. We employed a torsionally-constrained construct based on the same insert that was used to fabricate the Flap-DNA substrate. Before proceeding with measurements, force-extension curves of the tethers ensured they were completely single-stranded. The mechanical response of ssDNA with and without hRPA revealed an increase of extension, which was maximal at ∼3.8 pN (not shown). Further experiments confirmed that 20 nM RPA was saturating (**Figure S5C)**. At this applied force, the binding of 20 nM RPA produced an extraordinary increase of extension of the ssDNA tethers of ∼4 fold (**Figure 5E**). Next, while maintaining the RPA concentration, we introduced 5 nM HELB and 2 mM ATP into the fluid cell and recorded the effect of HELB activity. In half of the time-courses under these conditions (7 of 15 time-courses), the HELB and ATP caused loss of the beads (**Figure 5E**, black trace), probably because the translocating motor can disrupt the biotin-streptavidin bond holding the DNA to the bead. In the other half of the traces, we observed repetitive decreases in the bead’s height followed by a sudden recovery of the initial position (**Figure 5E**, red trace, **Figure 5F, and Figure S5E**). The reduction of the extension could be caused by the removal of RPA by HELB, as bare ssDNA has a shorter extension than the RPA filament. However, we can discard this possibility because free RPA is always present during the observed activity and would rebind the substrate on a faster timescale than HELB translocation, as clearly shown by the rapid extension rates observed upon initial RPA binding to ssDNA (**Figure 5E**). Instead, we interpret the data in terms of a looping model, in which the HELB remains fixed to one strand while reeling in downstream ssDNA via the motor domains (which may or may not be coupled to the clearance of RPA). In this scenario, the extension change rates we observe at different RPA concentrations are very similar to the translocation values measured in C-trap experiments, and the sudden recovery in height is simply explained as dissociation of HELB from DNA and release of the loop (compare rates in **Figure S5D** and **Figure 2D**). In complete contrast to these experiments with human RPA, no activity was observed in equivalent ssDNA MT experiments in the presence of 20 nM, 100 nM and 500 nM yeast RPA (**Figure S5F**, and not shown). Together these MT data suggest the possibility that HELB possesses a secondary static DNA binding site that facilitates the production of loops on RPA-coated ssDNA and further confirms the species specificity of the HELB-RPA interaction.

Intriguingly, bulk unwinding assays revealed a clear inhibitory effect on HELB helicase activity by RPA and SSB (**Figure S5G**). This was confirmed using MT-based unwinding assays, where we found that RPA *always* inhibited DNA helicase activity in Flap-DNA regardless of the applied force and whether the RPA was human or yeast in origin (not shown). A simple explanation could be that RPA or SSB prevent HELB loading to the free 5′ tail, but in any case, these results again focus our attention on *RPA-coated single-stranded DNA* as the physiological substrate for HELB and question the relevance of any DNA unwinding activity in its cellular function.

### Direct observation of HELB translocation on ssDNA shows that it is facilitated by, and causes displacement of, human RPA

We next sought to directly characterize how HELB interacts with and affects RPA filaments using combined optical trapping with confocal scanning microscopy. We labelled human RPA with the MB543 (RPA^MB543^) fluorophore (emission 570 nm) and used the dual-color imaging ability of our instrument to simultaneously detect HELB-QD (blue channel) and RPA^MB543^ (green channel) (**Figure 6A**). We first produced a ssDNA tether from lambda DNA as before. Then, we moved to a channel with 15 nM RPA^MB543^ and took a single image to confirm uniform coverage of the DNA by RPA (**Figure 6B**, upper green). We next moved the nucleoprotein complex to a channel containing 5 nM HELB-QD and 2 mM ATP (and no RPA), and acquired video scans and kymographs between the two beads. **Figure 6B** shows snapshots of a 2D video scan (**Supplementary Video 1**) where the movement of HELB-QDs (represented as blue dots) on an RPA^MB543^-ssDNA filament (represented in green) is observed. **Figure 6C** shows representative kymographs of the blue and green light excitation channels. We observed HELB trajectories in the blue channel moving unidirectionally, as observed previously on bare ssDNA (**Figure 2E**). The effect of HELB translocation on RPA distribution along the single strand of DNA was detected in the green channel. Interestingly, this was manifested as the progressive and unidirectional expansion of dark regions which were apparently cleared of RPA. The overlap of the blue and green emissions confirmed that the HELB trajectories match with the progression of the dark front (**Figure 6D** and **Figure S6A**). Notice, however, that in some dark regions of the RPA distribution we did not detect any correlated QD emission signal (see, for example, the red arrows in the merged kymograph of **Figure S6A**). We attribute this to the presence of untagged HELB proteins that could not be detected. As expected, in the absence of ATP we observed the binding of RPA and HELB, but the fluorescence signals remained static (**Figure S6B**). The translocation rate of HELB on RPA-covered ssDNA, deduced from the HELB-QDs trajectories (blue), was 66 ± 17 nt s^-1^ (peak ± w/2, R^2^ = 0.96, n = 23) (**Figure 6E**), considerably higher than on bare ssDNA.

**Figure 6.**
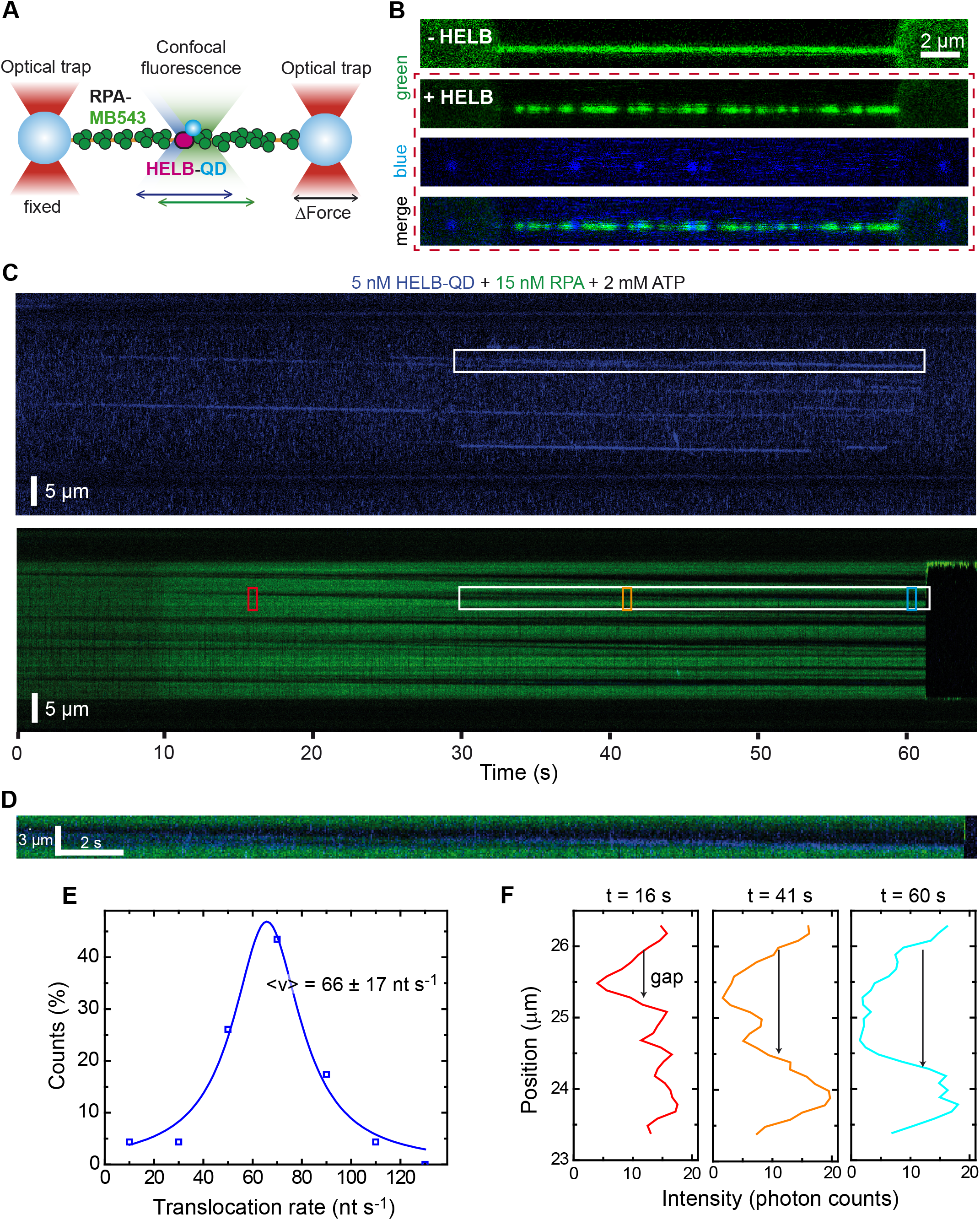
HELB translocation on ssDNA is facilitated by and causes displacement of human RPA. Illustration of the experimental C-trap setup. Individual ssDNA tethers attached to two optically-trapped beads are covered with fluorescent hRPA^MB543^ and exposed to HELB-QD. Both proteins were detected by two colour-excitation confocal microscopy. 2D scans of a tethered ssDNA covered by hRPA^MB534^ in the absence of HELB (top green) and after exposed to 5 nM HELB-QDs and 2 mM ATP filtered by green (RPA detection), blue (QDs detection) and merge fluorescence emission images. (**C**) Representative kymograph of HELB movement (blue) on ssDNA covered by hRPA^MB534^ (green) in the presence of 2 mM ATP. Signals obtained with blue and green emission filters are displayed separately. (**D**) Zoom area of the region marked in white in **C** which contains the merged fluorescence of the blue and green channels. (**E**) Distribution of HELB translocation rates on hRPA-covered ssDNA in the presence of 2 mM ATP. A Lorentzian fits the distribution with a peak at 66 ± 17 nt/s (n = 23). (**F**) Mean intensity profiles at different times along a trajectory of HELB on hRPA-covered ssDNA. Each line corresponds to the average intensity of 20 frames (0.82 s) that approximately comprised the corresponding coloured rectangles on the green kymograph in **C**. Experiments were done at forces between 14 and 23 pN.

Analysis of the human RPA distribution intensity profiles as a function of time suggests that HELB translocation both pushes RPA forward along the DNA and removes it from DNA (**Figure 6F**). We often observed an increase in the fluorescence intensity of RPA immediately in front of HELB, as if RPA accumulates ahead of the enzyme (compare, for example, red and orange lines, in **Figure 6F**). However, comparison of the intensity profiles between longer times also provides evidence for the removal of RPA from DNA: in some cases, we observed an increase in the RPA-free gap created by HELB without a compensatory increase in the intensity at the borders of the gap (compare red and cyan lines, in **Figure 6F**, for instance).

Equivalent experiments using fluorescent yeast RPA (yRPA^CY3^) gave strikingly different results. We observed that HELB binds poorly to ssDNA molecules covered by yRPA and the reduced number of HELB proteins bound exhibit very limited movement (**Figure S6C)**. Accordingly, the translocation rate distribution displayed a main peak around zero (**Figure S6D)**. As we showed no interaction between yRPA and HELB (**Figure 5B**), it is likely that the binding of the protein occurs in naked ssDNA regions, but HELB is blocked by yRPA and remains still. However, during the acquisition of long kymographs, we could detect movement of HELB, coupled with the clearance of yRPA from ssDNA. HELB moved in this case at 67 ± 30 nt s^-1^, a similar value than that observed in the presence of hRPA (**Figure 6E)**. Together these results are consistent with our previous findings showing that yeast RPA is inhibitory of the translocase activity of HELB.

## DISCUSSION

In this work we purified and characterised the human HELB helicase. This study confirms that the purified protein alone is an efficient ATPase and 5′ to 3′ ssDNA translocase with a coupling efficiency of 1 ATP per base. These properties are as expected based upon the similarity of the HELB helicase domain to the RecD-like family of SF1B helicases. We also demonstrated that ssDNA translocation is accompanied by the formation of DNA loops. Given that the protein is monomeric, this implies the presence of an additional static DNA binding domain beyond the translocating ssDNA binding site that is expected to be associated with the helicase core. The large binding site size observed here (∼20 bases) is also consistent with an additional unknown binding locus, because Superfamily 1 helicase domains bind a stretch of only about 8 bases (Gilhooly et al., 2013). DNA looping is an emerging feature of processive helicases which might help to supress reannealing and therefore improve duplex unwinding. However, HELB is in fact a very poor DNA helicase *in vitro*, and a cryptic ability to separate duplexes efficiently is only revealed by the application of an assisting force.

HELB was already shown to interact with the heterotrimeric RPA protein which plays a central and ubiquitous role in DNA replication and repair in eukaryotic cells (Caldwell and Spies, 2020; Sugitani and Chazin, 2015; Wold, 1997). Because of its high abundance and affinity for single-stranded DNA, RPA was originally thought of as a protective factor for single-stranded DNA. However, it is now appreciated that RPA nucleoprotein filaments can act as a dynamic platform for the recruitment or exclusion of other DNA binding proteins (Pokhrel et al., 2019), or for initiating cell signalling cues (Maréchal and Zou, 2015). Moreover, the ability of RPA to “melt out” secondary or alternative structures in DNA such as G-quadruplexes can also assist downstream processing of ssDNA intermediates, for example the formation of uniform RAD51 filaments to promote DNA strand exchange. Nevertheless, these useful roles of RPA in managing ssDNA intermediates present a paradox: how can a very tightly bound ssDNA be handed-off to additional enzymes that are required to complete replication or repair pathways? Interestingly, the RPA heterotrimer comprises six DNA binding domains joined by flexible linkers, and this modular organisation potentially allows for other proteins to bypass or access single-stranded DNA within RPA filaments despite the very high affinity interaction (Caldwell and Spies, 2020; Pokhrel et al., 2019). Such transactions may also be regulated by post-translational modifications to RPA, the use of alternative RPA subunits, or the targeted remodelling of RPA filaments by additional factors (Yates et al., 2018). Interestingly, RPA filaments are known to interact physically and functionally with several helicase or translocase enzymes (Awate and Brosh Jr, 2017). These interactions, typically with a basic patch in the RPA70 subunit, help recruit and activate the motor proteins to their physiological site of action, but may also have consequences for the formation, remodelling or removal of the RPA filament (Spies and Ha, 2010). Among the best studied examples are Superfamily 2 helicases such as WRN and HelQ, whose DNA unwinding activity is stimulated by the cognate RPA protein (Brosh et al., 1999; Jenkins et al., 2021). Similarly, the helicase activity of the archaeal nucleotide excision repair and transcription factor XPD is facilitated by RPA2 (Pugh et al., 2008). Remarkably, XPD can bypass RPA without displacing it, thereby overcoming its potential inhibitory effect as a roadblock to translocation (Honda et al., 2009).

Our work shows that the functional consequences of HELB-RPA interaction are profound for both proteins, but also quite distinctive compared to these other helicase systems. All ssDNA-dependent activities of HELB are stimulated by the presence of RPA. This phenomenon absolutely requires the cognate human RPA, and therefore we presume a sustained physical interaction, because the yeast orthologue and bacterial SSBs (data not shown) inhibit all activities of HELB under the same conditions. This latter observation makes good sense, because single-stranded DNA binding proteins interact tightly with nucleic acids and must therefore compete with HELB for access to ssDNA. Quantitative analysis of the single-stranded DNA dependent ATPase activity of HELB in the presence and absence of HELB provides support for the idea that RPA stimulates HELB both as the result of a recruitment/loading phenomenon and the activation of forward translocation, possibly because RPA pre-melts secondary structures which would otherwise slow the movement of the motor protein. Although it is both recruited and activated by its cognate RPA protein, HELB translocation activity then acts to remove RPA leaving naked ssDNA in its wake, and duplex unwinding is in fact inhibited by RPA. It seems clear therefore that the physiological target of HELB is an RPA-ssDNA nucleoprotein filament, and that HELB is not a classical helicase, but rather an RPA displacement motor, likely somewhat similar in function to the Srs2 helicase in yeast (Dhingra et al., 2021; De Tullio et al., 2017). These findings have important implications for better understanding the biochemical basis for the roles which HELB might play in DNA repair and replication.

HELB localizes to replication origins and appears to play a role in the onset of chromosomal replication via interactions with both CDC45, a component of the replicative helicase, and DNA polymerase α-primase, which synthesises RNA primers (Taneja et al., 2002). Successful firing of replication forks involves association of RPA with single-stranded DNA emerging from the CMG helicases as they undergo activation at S-phase, but RPA is also inhibitory to DNA polymerase α-primase (Bell and Labib, 2016; Taneja et al., 2002). Therefore, the RPA clearance activity of HELB observed here might facilitate the priming of DNA replication in the presence of the single-stranded DNA binding protein. HELB also localizes to chromatin in an RPA-dependent fashion during replication stress and might therefore also act during replication elongation in the recovery of stalled forks (Guler et al., 2012; Hazeslip et al., 2020). In this context, RPA clearance might facilitate the origin-independent assembly of the replisome or re-priming of leading strand replication.

Finally, HELB has been proposed to both enhance (Guler et al., 2012) and inhibit (Tkáč et al., 2016) homologous recombination by facilitating strand exchange and inhibiting DSB resection respectively. It is certainly easy to imagine how clearance of RPA in a 5′-to-3′ direction might promote strand exchange by facilitating the exchange of RPA for RAD51. Indeed, it is well-established that the inherent affinity of RAD51 for DNA is insufficient to compete effectively with RPA in the absence of mediator proteins such as BRCA2 (Jensen et al., 2010; Kowalczykowski, 2015). In contrast, it is less immediately obvious how the 5′-to-3′ translocation polarity of HELB could directly inhibit the DSB resection nucleases which generate 3′-ssDNA overhangs to initiate recombination. It is possible that removal of RPA might inhibit resection indirectly, since RPA has been shown to enhance this early processing step in DSB repair (Soniat et al., 2019). Alternatively, HELB might facilitate access to ssDNA for other factors that are inhibitory to DNA break resection and HR. Unpicking these possibilities will be the aim of future work.

## ACKNOWLEDGEMENTS

F.M.-H. acknowledges support from the European Research Council (ERC) under the European Union Horizon 2020 Research and Innovation Program (grant agreement 681299). Work in the Moreno-Herrero laboratory was also supported by Spanish Ministry of Economy and Competitiveness grant BFU2017-83794-P (AEI/FEDER, UE to F.M.-H.) and Comunidad de Madrid grants Tec4-Bio – S2018/NMT-4443 and NanoBioCancer

– Y2018/BIO-4747 (to F.M.-H.). Work in the Dillingham laboratory was supported by a Wellcome Trust Investigator Grant (100401/Z/12/Z to M.S.D.) and an Elizabeth Blackwell Early Career Fellowship from the University of Bristol (to O.J.W.). Work in the Antony laboratory was supported by grants (GM130746 and GM133967 to E.A.) from the National Institutes of Health.

Unlabelled yeast (*S. cerevisiae*) RPA was a kind gift from Luke Yates, and BirA was a kind gift from Charles Grummit.

## AUTHOR CONTRIBUTIONS

S.H. performed single molecule C-trap and magnetic tweezers experiments and analysis of the data. O.J.W. expressed and purified human HELB protein and performed molecular biology and biochemical characterization experiments and analysis of the data. C.A.-R. produced DNA substrates for single-molecule experiments. S.K and E.A. produced yeast RPA and human RPA proteins. F.M.-H. and S.H. designed the single-molecule experiments. M.S.D and O.J.W. designed the molecular biology and biochemical characterization experiments. M.S.D. and O.J.W. wrote the first draft of the manuscript with input from all authors. All authors critically reviewed the manuscript and approved the final version. M.S.D and F.M.-H. supervised the project.

## MATERIALS AND METHODS

### Human HELB protein expression and purification

A synthetic gene codon-optimised for *S. frugiperda* (Geneart, Invitrogen) encoding wild type human HELB was cloned into the pACEBac1 vector using the BamHI and XbaI restriction sites for use in the MultiBac system (Geneva Biotech). This was then screened for expression and purification using affinity tags in different positions. Following cleavage of a construct with a C-terminal 3C-cleavable StrepII tag, we obtained HELB in good yield and purity. The full-length recombinant protein as prepared (i.e. post-3C cleavage) contains a C-terminal extension-SGLEVLFQ (MW = 124,126 Da monomer), with the rest of the protein identical to UniProt entry Q8NG08. Mutagenesis (Quikchange XL, Agilent) was performed using this construct to create the K481A (ATPase) mutant. Bacmids were prepared by transposition of these plasmids and were used to transfect Sf9 cells in Insect Express media (Lonza) before viral amplification in the same cell line using standard techniques. For large scale expression, 500 mL of Hi5 cells at density 2 × 10^6^/mL were infected with 25 mL of P3 virus and harvested by centrifugation after 70 hours at 27°C with shaking. The pellets were lysed into buffer containing 50 mM Tris pH 8.0, 100 mM NaCl, 1 mM DTT, 10% glycerol, protease inhibitor cocktail (Roche) and then sonicated on ice for a total of 2 minutes. After centrifugation at 4°C for 30 mins at 50000g, the cleared lysate was applied to Streptactin beads (GE Healthcare) in batch and incubated for 1 h at 4°C with rotation. After washing five times in batch with buffer containing 20 mM Tris HCl pH 8.0, 100 mM NaCl, 5% glycerol, 1 mM DTT, the protein was then eluted in the same buffer containing 2.5 mM desthiobiotin. The HELB-containing fractions were then applied to a 5 mL Heparin column (GE Healthcare). After washing, HELB was eluted with a gradient from 100 mM to 1 M NaCl over 16 CV in buffer containing 20 mM Tris HCl pH 8.0, 5% glycerol, 1 mM DTT. The HELB-containing fractions were pooled and digested for 3h at 4°C with 3C protease to remove the StrepII tag. The cleavage reaction was run over a 5 mL Streptactin column (Qiagen) to remove any uncleaved HELB and free StrepII peptide and the cleaved HELB-containing flow-through collected. This was spin concentrated (50,000 Da cutoff, Millipore) down to a volume of approximately 200 µL and applied to a Superose6 10/300 column in buffer containing 20 mM Tris HCl pH 8.0, 200 mM NaCl, 5% glycerol, 1 mM DTT. The HELB peak eluted after a volume of 15.5 mL, and was then concentrated using a centrifugal filter unit (Millipore) with glycerol added to a final concentration of 10% (v/v) and the protein stored at −80°C. Protein concentration was determined using a theoretical extinction coefficient of 120,780 M^-1^ cm^-1^. Analysis of the purified protein by Orbitrap LC-MS/MS spectrometry was performed by the University of Bristol Mass Spectrometry Facility.

To produce biotinylated HELB, a modified version of the WT HELB pACEBac1 plasmid was constructed containing a C-terminal Avi-tag upstream of the cleavable StrepII tag. This protein expressed and purified in an identical manner to WT. After StrepII-tag cleavage and subsequent concentration to 50 µM, HELB-Avi was treated with BirA and biotin in order to site-specifically biotinylate the protein. After SEC purification (carried out as before) the protein was concentrated and then tested for biotinylation efficiency (measured as ∼100%). Conjugation of the biotinylated HELB protein to streptavidin did not affect its biochemical activities.

### Human and yeast RPA protein expression and purification

Human RPA was produced using plasmid p11d-hRPA (a kind gift from Mark Wold, Univ. of Iowa) and purified from *E. coli* as described (Binz et al., 2006). Fluorescent *S. cerevisiae* RPA was purified and labeled with Cy3 as described previously using non-canonical amino acids (Pokhrel et al NAR 2017, Pokhrel et al NSMB 2019). Fluorescent human RPA labeled with MB543 was also generated using non-canonical amino acids (ncAA) as described for yeast RPA (Kuppa et al., 2021; Pokhrel et al., 2017, 2019). Briefly, to obtain RPA carrying 4-aziodophenylalanine (4AZP), a p11d-hRPA-TAG32-107 plasmid was generated from p11d-hRPA. This plasmid contains a TAG at the chosen site of fluorophore incorporation, which corresponds to W107 in the human RPA32 subunit. A 6x-poly-histidine affinity tag was also engineered at the C-terminus of RPA32. This plasmid was cotransformed into BL21PlysS cells with pDule2-pCNF (Hammill et al., 2007). This plasmid codes for the orthogonal tRNA^UAG^ and tRNA synthetase for 4AZP incorporation. Cotransformants were selected using both ampicillin (100 μg/mL) and spectinomycin (50 μg/mL). An overnight culture (10 mL) from a single colony was grown in LB media containing ampicillin and spectinomycin. 10 mL of the overnight culture was added to 1 L of minimal media. The minimal media for ncAA incorporation was prepared as previously described (Hammill et al., 2007). Cells were grown at 37°C until the OD_600_ reached 2.0 and then induced with 0.4 mM IPTG along with 1 mM 4AZP. The 4AZP solution was prepared by first dissolving 206 mg in 250 μL of 5 M NaOH, vortexed extensively, and then adjusted to 8 ml with H_2_O, and the entire mixture was added to 1 L of media to achieve a final concentration of 1 mM. Induction was carried out at 37°C for 3 hours. Harvested cells were resuspended in 120 mL cell resuspension buffer (30 mM HEPES, pH 7.8, 300 mM KCl, 0.02% Tween-20, 1.5X protease inhibitor cocktail, 1 mM PMSF, 10% (v/v) glycerol and 10 mM imidazole). Cells were lysed using 400 mg/mL lysozyme followed by sonication. Clarified lysates were fractionated on a Ni^2+^-NTA agarose column. Protein was eluted using cell resuspension buffer containing 400 mM imidazole. Fractions containing RPA were pooled and diluted with H^0^ buffer (30 mM HEPES, pH 7.8, 300 mM KCl, 0.02% Tween-20, 1.5X protease inhibitor cocktail, 10% (v/v) glycerol and 25 mM EDTA pH 8.0) to match the conductivity of buffer H^100^, and further fractionated over a fast-flow Heparin column. RPA was eluted using a linear gradient H^100^–H^1500^, and fractions containing RPA were pooled and concentrated using an Amicon spin concentrator (30 kDa molecular weight cut-off). The concentrated RPA was next loaded onto a S200 column and fractionated using RPA storage buffer (30 mM HEPES, pH 7.8, 30 mM KCl, 0.25 mM EDTA, 0.02% Tween-20, and 10% (v/v) glycerol).

To fluorescently label RPA, RPA^AZP^ (∼ 4 µM in 5 mL of storage buffer) was mixed with 1.5-fold molar excess DBCO-MB543 (Click Chemistry Tools Inc., AZ) or DBCO-Cy3. The reaction was incubated for 2 hours at 4°C and the labeled RPA was separated from free dye on a Biogel P4 column and resolved using RPA-storage buffer. An additional 10% glycerol was added to the fluorescent RPA protein and flash frozen using liquid nitrogen and stored at –80°C. RPA concentration was measured spectroscopically using e_280_ = 98500 M^-1^cm^-1^ (yRPA) or 87,210 M^-1^cm^-1^ (hRPA) and corrected for fluorophore contributions as described (Kuppa et al., 2021).

### Electrophoretic Mobility Shift DNA binding assays

5′-Cy5-labelled DNA substrates (**Supplementary Table S2**) (2.5 nM final) were mixed with increasing amounts of HELB protein in a total volume of 10 μL 1X EMSA buffer (20 mM Tris HCl pH 8.0, 100 mM NaCl, 1 mM DTT, 0.1 mg/mL BSA, 5% glycerol) and then incubated for 10 min at 25°C. In the case where RPA was used, it was added to the DNA before incubation with HELB. The samples were then loaded onto a 6% polyacrylamide (29:1) native 1xTBE gel and separated by electrophoresis in 1xTBE at 150V for 40 mins. The gels were visualised using a Typhoon scanner and analysed using ImageQuant software.

### PIFE DNA binding assays

3′-Cy3-labelled DNA substrates (**Supplementary Table S2**) (2 nM final) were mixed in a quartz cuvette (Hellma) with increasing amounts of HELB protein in a total volume of 150 μL 1X PIFE buffer (20 mM Tris HCl pH 8.0, 100 mM NaCl, 1 mM DTT). After 1 min, the protein DNA mixture was scanned (λ_Ex_=530 nm, λ_Em_=562 nm, 5 nm slit widths) and the fluorescence intensity measured in arbitrary units. The fully saturated DNA-HELB gave an increase in fluorescence of around 60% compared to the original signal of free DNA. The measurements were normalised with the initial reading being 0 and the highest reading being 100%. Data was fitted to a weak binding isotherm *y* = *Bmax*. *x*/(*x* + *K*_*d*_) + *C*, where *K*_*d*_ is the binding constant, the protein concentration, and the substrate bound under saturating conditions.

For the binding stoichiometry plots, DNA was used at high concentration (stated in the plots) and the experiments were carried out as before. These data were fitted to a version of the tight binding equation

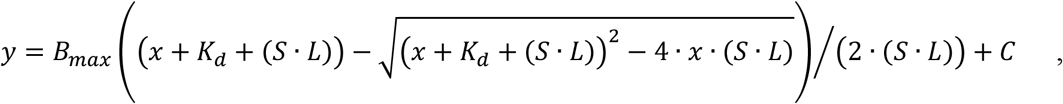

where *S* stands for stoichiometry (number of proteins bound to the DNA substrate), *L* is the DNA concentration, *K*_*d*_ is the binding constant, and *x* the protein concentration.

### ATPase assays

ATPase activity was measured by coupling the hydrolysis of ATP to the oxidation of NADH which gives a change in absorbance at 340 nm. Reactions were performed in a buffer containing 20 mM Tris-HCl pH 8.0, 50 mM NaCl, 5 mM DTT, 1 mM MgCl_2_, 50 U/mL lactate dehydrogenase, 50 U/mL pyruvate dehydrogenase, 1 mM PEP and 100 µg/mL NADH. Rates of ATP hydrolysis were measured over 1 min at 25°C. For calculation of K_DNA_, the ATP concentration was fixed at 4 mM and the Michaelis-Menten plot was performed at [Poly(dT)] =10 × K_DNA_. The concentration of HELB was 50 nM in these assays. In the cases where different DNA substrates and/or RPA were used, their concentrations are stated in the plots and/or figure legends.

### ssDNA translocation assays

Streptavidin displacement assays were based on the method of Morris and Raney (Morris and Raney, 1999) with minor modifications as in (Yeeles et al., 2011). 5 nM (molecules) of 5′-^32^P-labelled substrate 45mer oligonucleotides (**Supplementary Table S2**) were incubated with 400 nM streptavidin in 25 mM Tris-HCl pH 8, 50 mM NaCl, 4 mM MgCl_2,_ 1mM TCEP. Substrates were modified with either a 5′ or 3′ biotin moiety as indicated. The reaction was initiated by adding an equal volume of protein solution in the same buffer to give final concentrations of 250 nM HELB, 5 mM ATP and 8 µM biotin. The reaction was incubated at 37°C and stopped at certain points within a 10 min time course by quenching with an equal volume of stop buffer (300 mM EDTA, 400 mM NaCl, 30 µM poly(dT)). The products were separated on 10% polyacrylamide 1xTBE gels and visualised by phosphorimaging using a Typhoon imager. In the case where RPA was included, the reactions were set up as before but stopped after five minutes, and the final RPA concentrations used were 0, 25, 50, 100 and 200 nM.

### Helicase (strand displacement) assays

Strand-displacement assays were based on a modification of the method of Matson (Matson et al., 1983). 10 nM (molecules) of 5′-Cy5-labelled oligonucleotides (**Supplementary Table S2**)consisting of a 25 bp duplex region with a flanking ssDNA overhang of 20 nt were incubated with 500 nM HELB in 20 mM Tris-HCl pH 8.0, 4 mM MgCl_2_, 4 mM ATP, 1 mM DTT for 5 min at 25°C. The reaction was quenched by adding an equal volume of stop buffer (200 mM EDTA, 1% SDS, 10% (w/v) Ficoll 400 and 100 nM of an unlabelled form of the labelled strand in the substrate. In the cases where RPA was included, the final concentration was 100 nM. The products were separated on 15% polyacrylamide 1xTBE gels and visualised using a Typhoon imager.

### Blue Native PAGE

Proteins and DNA were mixed in equimolar amounts, in 20 mM Tris pH 8.0, 200 mM NaCl, 1 mM DTT and after 5 min incubation at room temperature samples were mixed with sample buffer and loaded onto a pre-cast blue native gel and run according to the manufacturer’s instructions (Life).

### Analytical SEC and SEC-MALS

Proteins and DNA were mixed in equimolar amounts and loaded onto a Superose6 10/300 column equilibrated in 20 mM Tris pH 8.0, 200 mM NaCl, 1 mM TCEP. Chromatograms were recorded for the absorbance at 280 nm (Abs_280_) and at 260 nm (Abs_260_) against volume (mL) using Unicorn. SEC-MALS was used to determine the absolute molecular masses of full-length HELB. A 50 µg sample of HELB was loaded at 0.5 mL/min onto a Superose 6 10/300 size-exclusion chromatography column (GE Healthcare) in 20 mM Tris pH 8.0, 200 mM NaCl, 1 mM TCEP using an Agilent HPLC. The eluate from the column was coupled to a DAWN HELEOS II MALS detector (Wyatt Technology) and an Optilab T-rEX differential refractometer (Wyatt Technology). ASTRA 6 software (Wyatt Technology) was used to collect and analyse light scattering and differential refractive index data according to the manufacturer’s instructions. Molecular mass and estimated error were calculated across individual eluted peaks.

### Optical tweezers and confocal microscopy

Correlative tweezers-fluorescence experiments were performed at room temperature on an instrument combining three-color confocal fluorescence microscopy with dual trap optical tweezers (C-Trap, LUMICKS). A computer-controlled stage enabled the fast displacement of the optical traps within a five-channel fluid cell (**Figure S2A**). This microfluidic cell allowed for the *in-situ* assembly and characterization of dumbbell DNA constructs, and facilitated the direct transfer of the tethered DNA between different flow channels. Laminar flow separated channels 1-3 were used to form a single-stranded biotin-λ DNA tether as follows: a single 4.38-µm streptavidin-coated polystyrene bead (Spherotech) was caught in each trap in channel 1 (trap stiffness of ∼0.4 pN/nm). The traps were then moved to channel 2 containing the biotinylated DNA intended for ssDNA fabrication by force. This DNA is a λ-phage dsDNA (LUMICKS) biotinylated at the 3′ and 5′ positions of the same strand so that the non-biotinylated strand can be removed by tension (Candelli et al., 2013; Wasserman et al., 2019). The traps were then moved to channel 3 containing a low ionic strength buffer (10 mM Tris pH 8.0, 1 mM EDTA) to favour the peeling. Here the duplex DNA was held at forces higher than the overstretching transition and the non-biotinylated strand was removed (**Figure S2B**). The presence of a single-stranded DNA was verified by force-extension curves (**Figure S2C**). Orthogonal channels 4 and 5 were used for protein loading and imaging.

To investigate the ability of HELB to bind and translocate on ssDNA, the tethers were moved into channel 5 containing 5 nM biotinylated HELB-QDs in 20 mM Tris pH 7.5, 30 mM NaCl, 4 mM MgCl_2_, 5 mM DTT (reaction buffer) and 2 mM ATP. To test the behaviour of HELB on a ssDNA covered by RPA, the single-stranded tethers were first moved to channel 4 containing 15 nM human RPA^MB543^ or *S. cerevisiae* RPA^CY3^ in reaction buffer. The RPA-loaded tether was then dragged to channel 5 containing 5 nM HELB-QDs in the reaction buffer supplemented with 2 mM ATP. Flow was turned off during data acquisition. The instrument is equipped with a multi wavelength CW laser engine for 3 color confocal imaging. For our experiments, two excitation lasers were used, 488 nm for QDs-525 and 532 nm for MB543 and Cy3 fluorophores. The emission was detected employing a blue filter 512/25 nm and a green filter 585/75 nm. Kymographs were generated via a confocal line scan through the centre of the two beads.

Force and fluorescence data were analyzed using FIJI (Schindelin et al., 2012) and custom software provided by LUMICKS. HELB velocity on bare ssDNA and in the presence of RPA was measured by dividing the distance travelled by the duration of each trajectory, ΔL/Δt (**Figure S6A**). The HELB-QD trajectories were analysed from the signal detected in the blue emission filter 512/25 nm.

### Conjugation of HELB with quantum dots

To label HELB with QDs we incubated a 1:5 molar ratio solution of biotinylated HELB and streptavidin-coated QDs 525 (Q10143, Invitrogen) for 30 min on ice. We then added 1 mM biotin (B4501, Sigma) to neutralize the streptavidin molecules not bound to HELB. After 10 min the mixture was diluted in the reaction buffer to a final concentration of 5 nM HELB-QD and readily used.

### DNA substrates for Magnetic Tweezers experiments

The Flap-DNA substrate for magnetic tweezers contains a 5′-ssDNA overhang or flap sequence of 37 poly(dT) nucleotides at a specific-site. It consists of a central fragment ligated to two digoxigenin or biotin-labelled DNA handles. The substrate is based on the pNLrep plasmid (Luzzietti et al., 2011) that has a DNA sequence with five closely-spaced BbvCI restriction sites. Nicking of one of the two strands with Nt.BbvCI enzyme results in the formation of short 15–16 bases long fragments after heat denaturation, leaving a 63 nucleotides gap, where desired oligonucleotides can be hybridised.

The Flap-DNA substrate was fabricated following the protocol described in (Wilkinson et al., 2020) with slight modifications. The pNLrep plasmid was digested with KpnI and PsiI enzymes (NEB) giving a 6337 bp product. To avoid reannealing of Nt.BbvCI cleavage products, a 100X excess of short oligonucleotides complementary to the four released Nt.BbvCI-fragments (**Supplementary Table S1**) were added after Nt.BbvCI digestion and before inactivation of the enzyme for 20 min at 80°C. Following enzyme inactivation, the sample was cooled down to 40°C at a 1°C/1 min rate. The 6337 bp product with a 63 nt-gap was then purified with a PCR purification kit from QIAGEN and a 150X excess of Poly(dT)-flap oligonucleotide (**Supplementary Table S1**) was hybridised as described in (Wilkinson et al., 2020). This generated a substrate as shown in (**Figure 3B**) with a duplex region of 4563 bp from the base of the Poly(dT)-tail oligonucleotide to the magnetic bead and a 37 nt poly(dT) tail. Handles were PCR-generated from the plasmid pSP73-JY0 (Fili et al., 2010) using appropriate oligonucleotides (**Supplementary Table S1**) and Bio-dUTP or Dig-dUTP (Roche), followed by restriction with PsiI or KpnI. The labelled fragments were ligated to the central part and the excess of oligonucleotides removed using two Microspin S-400 columns. DNAs were never exposed to intercalant dyes or UV radiation during their production and were stored at 4°C.

The Gap-DNA substrate was prepared as the Flap-DNA substrate but omitting the steps of hybridisation of the Poly(dT)-tail oligonucleotide after digestion with Nt.BbvCI.

A torsionally-constrained DNA substrate was prepared as the Flap-DNA substrate but omitting the steps of digestion with Nt.BbvCI to create the gap. A small fraction of these molecules were nevertheless nicked and were used in control experiments as Nicked DNA substrates.

### Magnetic tweezers assays

We used a magnetic tweezers instrument setup similar to the one reported previously (Carrasco et al., 2013). Raw data was recorded at 120 Hz and filtered to 6 Hz for representation. Force values were calculated using the Brownian motion method applied to a DNA-tethered bead (Strick et al., 1998). Flap-DNA substrate (**Figure 3B**) essentially consists of a DNA molecule of ∼6.3 kbp containing a flap poly(dT) tail of 37 nt in a specific-site, and flanked by two smaller fragments (∼1 kb) that act as the immobilization handles as they are labelled with biotins or digoxigenins. The labelled parts are used to specifically bind each DNA end to a glass surface covered by anti-digoxigenins and to streptavidin coated magnetic beads. A control Gap-DNA substrate (with the gap but without a flap sequence) (**Figure S2F**) and a torsionally-constrained dsDNA substrate (without a gap, employed to get ssDNA tethers) were also prepared. Doubly-tethered beads were identified by applying magnet rotations on the beads and not considered for the analysis. A detailed description of the construction of the magnetic tweezers DNA substrates can be found in the **Supplementary Methods**. DNA oligonucleotides used to fabricate magnetic tweezers substrates can be found in **Supplementary Table S1**. Sequence of the DNA fragment used in this work can be found in **Supplementary Table S3**.

Single-molecule unwinding experiments were carried out at ambient temperature and at 1 or 8.4 pN as indicated, in a reaction buffer that contained 20 mM Tris-HCl pH 7.5, 30 mM NaCl, 4 mM MgCl_2_, 5 mM DTT. To initiate the reaction, 100 nM HELB and 1 mM ATP were flowed into the fluid cell at 20 µL/min while the positions of the beads were measured by video microscopy. The injection of proteins is indicated as shadowed regions in the time courses. A fluid cell made with two parafilm layers (100 µL volume) and vertical alignment magnets with 0.2 mm gap were used to reach high applied forces using 1 µm beads (Dynabeads, Invitrogen). We chose to work at this concentration of HELB because we detected very limited activities below 100 nM, insufficient to manage a proper statistical analysis.

To produce single-stranded DNA molecules for MT experiments we followed a protocol previously described in (Carrasco et al., 2020). Briefly, it consists in heating at 95°C for 5 minutes the stock solution of DNA in buffer TD (10 mM Tris-HCl pH 8, 20 mM DTT) followed by fast cooling to 4°C. For this purpose, we employed a torsionally-constrained DNA construct based on the same insert that was used to fabricate the DNA with a flap. In this case, the DNA was not treated with nucleases to generate any gap so that the molecule had no nick and was labelled with digoxigenins and biotins in both strands of the corresponding handles. The separation of the two strands by heat provided us with two single-stranded DNA molecules which can bind the surface and the bead in our assay. Note that we have the same probability of finding the biotinylated handle at 3′-end and at the 5′-end. The ssDNA molecules were then introduced into the fluid cell in ice-chilled TD buffer and incubated for several minutes. Then, we flowed in 2.8 µm paramagnetic beads covered by streptavidin (M280, Invitrogen) and incubated them briefly. After washing to remove the non-bound molecules and beads, we applied force with vertical alignment magnets with a 1 mm gap to locate the ssDNA tethers.

The quoted distances in base pairs were calculated from changes in DNA extension considering the different stretching properties of ssDNA and dsDNA. We experimentally determined the mechanical properties of dsDNA and ssDNA from force-extension curves in HELB’s reaction buffer. For dsDNA we use the value given by the worm-like chain model of rise per base pair at a given force. For ssDNA, fits to the Freely Jointed Chain model at forces larger than 8 pN, where the formation of ssDNA secondary structure is prevented by force, resulted in a contour length per base of 0.66 nm nt^-1^, in agreement with previously reported values (Bosco et al., 2014).

## REFERENCES

Awate, S., and Brosh Jr, R.M. (2017). Interactive Roles of DNA Helicases and Translocases with the Single-Stranded DNA Binding Protein RPA in Nucleic Acid Metabolism. Int. J. Mol. Sci. 18, 1233.

Bell, S.P., and Labib, K. (2016). Chromosome Duplication in Saccharomyces cerevisiae. Genetics 203, 1027–1067.

Bosco, A., Camunas-Soler, J., and Ritort, F. (2014). Elastic properties and secondary structure formation of single-stranded DNA at monovalent and divalent salt conditions. Nucleic Acids Res 42, 2064–2074.

Brosh, R.M.J., Orren, D.K., Nehlin, J.O., Ravn, P.H., Kenny, M.K., Machwe, A., and Bohr, V.A. (1999). Functional and physical interaction between WRN helicase and human replication protein A. J. Biol. Chem. 274, 18341–18350.

Bustamante, C., Bryant, Z., and Smith, S.B. (2003). Ten years of tension: single-molecule DNA mechanics. Nature 421, 423–427.

Caldwell, C.C., and Spies, M. (2020). Dynamic elements of replication protein A at the crossroads of DNA replication, recombination, and repair. Crit. Rev. Biochem. Mol. Biol. 55, 482–507.

Candelli, A., Wuite, G.J., and Peterman, E.J. (2011). Combining optical trapping, fluorescence microscopy and micro-fluidics for single molecule studies of DNA-protein interactions. Phys Chem Chem Phys 13, 7263–7272.

Candelli, A., Hoekstra, T.P., Farge, G., Gross, P., Peterman, E.J.G., and Wuite, G.J.L. (2013). A toolbox for generating single-stranded DNA in optical tweezers experiments. Biopolymers 99, 611–620.

Carrasco, C., Gilhooly, N.S., Dillingham, M.S., and Moreno-Herrero, F. (2013). On the mechanism of recombination hotspot scanning during double-stranded DNA break resection. Proc. Natl. Acad. Sci. U. S. A. 110, E2562–E2571.

Carrasco, C., Pastrana, C.L., Aicart-Ramos, C., Leuba, S.H., Khan, S.A., and Moreno-Herrero, F. (2020). Dynamics of DNA nicking and unwinding by the RepC-PcrA complex. Nucleic Acids Res. 48, 2013–2025.

Day, F.R., Ruth, K.S., Thompson, D.J., Lunetta, K.L., Pervjakova, N., Chasman, D.I., Stolk, L., Finucane, H.K., Sulem, P., Bulik-Sullivan, B., et al. (2015). Large-scale genomic analyses link reproductive aging to hypothalamic signaling, breast cancer susceptibility and BRCA1-mediated DNA repair. Nat. Genet. 47, 1294–1303.

Dessinges, M.N., Lionnet, T., Xi, X.G., Bensimon, D., and Croquette, V. (2004). Single-molecule assay reveals strand switching and enhanced processivity of UvrD. Proc Natl Acad Sci U S A 101, 6439–6444.

Dhingra, N., Kuppa, S., Wei, L., Pokhrel, N., Baburyan, S., Meng, X., Antony, E., and Zhao, X. (2021). The Srs2 helicase dampens DNA damage checkpoint by recycling RPA from chromatin. Proc. Natl. Acad. Sci. U. S. A. 118, e2020185118.

Dueva, R., and Iliakis, G. (2020). Replication protein A: a multifunctional protein with roles in DNA replication, repair and beyond. NAR Cancer 2.

Gerhardt, J., Guler, G.D., and Fanning, E. (2015). Human DNA helicase B interacts with the replication initiation protein Cdc45 and facilitates Cdc45 binding onto chromatin. Exp. Cell Res. 334, 283–293.

Gilhooly, N.S., Gwynn, E.J., and Dillingham, M.S. (2013). Superfamily 1 helicases. Front. Biosci. (Schol. Ed). 5, 206–216.

Guler, G.D., Liu, H., Vaithiyalingam, S., Arnett, D.R., Kremmer, E., Chazin, W.J., and Fanning, E. (2012). Human DNA helicase B (HDHB) binds to replication protein A and facilitates cellular recovery from replication stress. J. Biol. Chem. 287, 6469–6481.

Guler, G.D., Rosenwaks, Z., and Gerhardt, J. (2018). Human DNA Helicase B as a Candidate for Unwinding Secondary CGG Repeat Structures at the Fragile X Mental Retardation Gene. Front. Mol. Neurosci. 11, 138.

Hammill, J.T., Miyake-Stoner, S., Hazen, J.L., Jackson, J.C., and Mehl, R.A. (2007). Preparation of site-specifically labeled fluorinated proteins for 19F-NMR structural characterization. Nat. Protoc. 2, 2601–2607.

Hazeslip, L., Zafar, M.K., Chauhan, M.Z., and Byrd, A.K. (2020). Genome Maintenance by DNA Helicase B. Genes (Basel). 11, 578.

Honda, M., Park, J., Pugh, R.A., Ha, T., and Spies, M. (2009). Single-molecule analysis reveals differential effect of ssDNA-binding proteins on DNA translocation by XPD helicase. Mol. Cell 35, 694–703.

Jenkins, T., Northall, S.J., Ptchelkine, D., Lever, R., Cubbon, A., Betts, H., Taresco, V., Cooper, C.D.O., McHugh, P.J., Soultanas, P., et al. (2021). The HelQ human DNA repair helicase utilizes a PWI-like domain for DNA loading through interaction with RPA, triggering DNA unwinding by the HelQ helicase core. NAR Cancer 3.

Jensen, R.B., Carreira, A., and Kowalczykowski, S.C. (2010). Purified human BRCA2 stimulates RAD51-mediated recombination. Nature 467, 678–683.

Korolev, S., Hsieh, J., Gauss, G.H., Lohman, T.M., and Waksman, G. (1997). Major domain swiveling revealed by the crystal structures of complexes of E. coli Rep helicase bound to single-stranded DNA and ADP. Cell 90, 635–647.

Kowalczykowski, S.C. (2015). An Overview of the Molecular Mechanisms of Recombinational DNA Repair. Cold Spring Harb Perspect Biol 7, a016410.

Kuppa, S., Pokhrel, N., Corless, E., Origanti, S., and Antony, E. (2021). Generation of Fluorescent Versions of Saccharomyces cerevisiae RPA to Study the Conformational Dynamics of Its ssDNA-Binding Domains. Methods Mol. Biol. 2281, 151–168.

Liu, H., Yan, P., and Fanning, E. (2015). Human DNA helicase B functions in cellular homologous recombination and stimulates Rad51-mediated 5’-3’ heteroduplex extension in vitro. PLoS One 10, e0116852.

Maréchal, A., and Zou, L. (2015). RPA-coated single-stranded DNA as a platform for post-translational modifications in the DNA damage response. Cell Res. 25, 9–23.

Matson, S.W., Tabor, S., and Richardson, C.C. (1983). The gene 4 protein of bacteriophage T7. Characterization of helicase activity. J. Biol. Chem. 258, 14017–14024.

Morris, P.D., and Raney, K.D. (1999). DNA helicases displace streptavidin from biotin-labeled oligonucleotides. Biochemistry 38, 5164–5171.

Newton, M.D., Taylor, B.J., Driessen, R.P.C., Roos, L., Cvetesic, N., Allyjaun, S., Lenhard, B., Cuomo, M.E., and Rueda, D.S. (2019). DNA stretching induces Cas9 off-target activity. Nat. Struct. Mol. Biol. 26, 185–192.

Pincus, D.L., Chakrabarti, S., and Thirumalai, D. (2015). Helicase Processivity and Not the Unwinding Velocity Exhibits Universal Increase with Force. Biophys. J. 109, 220–230.

Pokhrel, N., Origanti, S., Davenport, E.P., Gandhi, D., Kaniecki, K., Mehl, R.A., Greene, E.C., Dockendorff, C., and Antony, E. (2017). Monitoring Replication Protein A (RPA) dynamics in homologous recombination through site-specific incorporation of non-canonical amino acids. Nucleic Acids Res. 45, 9413–9426.

Pokhrel, N., Caldwell, C.C., Corless, E.I., Tillison, E.A., Tibbs, J., Jocic, N., Tabei, S.M.A., Wold, M.S., Spies, M., and Antony, E. (2019). Dynamics and selective remodeling of the DNA-binding domains of RPA. Nat. Struct. Mol. Biol. 26, 129–136.

Pugh, R.A., Lin, Y., Eller, C., Leesley, H., Cann, I.K.O., and Spies, M. (2008). Ferroplasma acidarmanus RPA2 facilitates efficient unwinding of forked DNA substrates by monomers of FacXPD helicase. J. Mol. Biol. 383, 982–998.

Saikrishnan, K., Griffiths, S.P., Cook, N., Court, R., and Wigley, D.B. (2008). DNA binding to RecD: role of the 1B domain in SF1B helicase activity. EMBO J. 27, 2222–2229.

Saitoh, A., Tada, S., Katada, T., and Enomoto, T. (1995). Stimulation of mouse DNA primase-catalyzed oligoribonucleotide synthesis by mouse DNA helicase B. Nucleic Acids Res. 23, 2014–2018.

Schindelin, J., Arganda-Carreras, I., Frise, E., Kaynig, V., Longair, M., Pietzsch, T., Preibisch, S., Rueden, C., Saalfeld, S., Schmid, B., et al. (2012). Fiji: An open-source platform for biological-image analysis. Nat. Methods 9, 676–682.

Soniat, M.M., Myler, L.R., Kuo, H.-C., Paull, T.T., and Finkelstein, I.J. (2019). RPA Phosphorylation Inhibits DNA Resection. Mol. Cell 75, 145-153.e5.

Spies, M., and Ha, T. (2010). Inching over hurdles: How DNA helicases move on crowded lattices. Cell Cycle 9, 1742–1749.

Strick, T.R., Allemand, J.F., Bensimon, D., and Croquette, V. (1998). Behavior of supercoiled DNA. Biophys. J. 74, 2016–2028.

Sugitani, N., and Chazin, W.J. (2015). Characteristics and concepts of dynamic hub proteins in DNA processing machinery from studies of RPA. Prog. Biophys. Mol. Biol. 117, 206–211.

Taneja, P., Gu, J., Peng, R., Carrick, R., Uchiumi, F., Ott, R.D., Gustafson, E., Podust, V.N., and Fanning, E. (2002). A dominant-negative mutant of human DNA helicase B blocks the onset of chromosomal DNA replication. J. Biol. Chem. 277, 40853–40861.

Tate, J.G., Bamford, S., Jubb, H.C., Sondka, Z., Beare, D.M., Bindal, N., Boutselakis, H., Cole, C.G., Creatore, C., Dawson, E., et al. (2019). COSMIC: the Catalogue Of Somatic Mutations In Cancer. Nucleic Acids Res. 47, D941–D947.

Tkác, J., Xu, G., Adhikary, H., Young, J.T.F., Gallo, D., Escribano-Díaz, C., Krietsch, J., Orthwein, A., Munro, M., Sol, W., et al. (2016). HELB Is a Feedback Inhibitor of DNA End Resection. Mol. Cell 61, 405–418.

De Tullio, L., Kaniecki, K., Kwon, Y., Crickard, J.B., Sung, P., and Greene, E.C. (2017). Yeast Srs2 Helicase Promotes Redistribution of Single-Stranded DNA-Bound RPA and Rad52 in Homologous Recombination Regulation. Cell Rep. 21, 570–577.

Wasserman, M.R., Schauer, G.D., O’Donnell, M.E., and Liu, S. (2019). Replication Fork Activation Is Enabled by a Single-Stranded DNA Gate in CMG Helicase. Cell 178, 600-611.e16.

Wilkinson, O.J., Carrasco, C., Aicart-Ramos, C., Moreno-Herrero, F., and Dillingham, M.S. (2020). Bulk and single-molecule analysis of a bacterial DNA2-like helicase-nuclease reveals a single-stranded DNA looping motor. Nucleic Acids Res. 48, 7991–8005.

Wold, M.S. (1997). Replication protein A: a heterotrimeric, single-stranded DNA-binding protein required for eukaryotic DNA metabolism. Annu. Rev. Biochem. 66, 61–92.

Yates, L.A., Aramayo, R.J., Pokhrel, N., Caldwell, C.C., Kaplan, J.A., Perera, R.L., Spies, M., Antony, E., and Zhang, X. (2018). A structural and dynamic model for the assembly of Replication Protein A on single-stranded DNA. Nat. Commun. 9, 5447.

Yeeles, J.T., Gwynn, E.J., Webb, M.R., and Dillingham, M.S. (2011). The AddAB helicase-nuclease catalyses rapid and processive DNA unwinding using a single Superfamily 1A motor domain. Nucleic Acids Res 39, 2271–2285.

Zou, Y., Liu, Y., Wu, X., and Shell, S.M. (2006). Functions of human replication protein A (RPA): from DNA replication to DNA damage and stress responses. J. Cell. Physiol. 208, 267–273.

