## Supplemental figures and captions for "Human HELB is a processive motor protein which catalyses RPA clearance from single-stranded DNA"

#### SUPPLEMENTARY FIGURE LEGENDS

##### Figure S1. HELB binding constant and binding stoichiometry determination

(**A**) PIFE DNA binding assay at low DNA concentration ($<K_D$) shows that tightest binding occurs with ssDNA of length >30 bases suggesting the presence of an additional DNA binding site in HELB other than that in the RecD-like domain. (**B-E**) PIFE DNA binding assays performed at high DNA concentrations ($>>K_D$) and fitted to the tight binding equation allow the binding stoichiometry to be determined for each length of DNA tested. Together, the data suggest a DNA-binding site size for HELB of approximately 20 nucleotides and is consistent with the presence of an additional DNA binding domain outside the RecD-like helicase domain.

##### Figure S2. Setup for optical tweezers experiments and ssDNA *in situ* formation

(**A**) Schematic of the experimental fluid cell used for optical tweezers experiments. Individual DNA tethers were formed in channels 1-3 separated by laminar flow containing streptavidin-coated beads, biotinylated λ-DNA for single-stranded DNA formation and a buffer with low salt, respectively. After ssDNA formation, the traps were subsequently moved to channels 4 and 5 for protein loading and imaging. (**B**) Schematic illustrating *in situ* formation of a ssDNA tether using a dual-trap optical tweezers. A single dsDNA biotinylated on both ends, but on the same strand, is captured with the optical traps. Then, the non-biotinylated strand is removed by force-induced melting of the duplex. (**C**) Force-extension curves indicate the transition from dsDNA (black) to ssDNA (red). (**D**) Example of a kymograph with HELB trajectories moving upwards. (**E**) Representative kymograph showing that HELB binds but remain stationary on bare ssDNA in the absence of ATP. (**F**) Scheme of the Gap-DNA substrate, a dsDNA molecule with a gap of 63 nt and no 5′-flap. (**G**) HELB requires a 5′-flap to unwind duplex DNA at low force. Representative time-courses of MT experiments with Gap-DNA and 100 nM HELB and 1 mM ATP at 1 pN, show no HELB unwinding activity.

##### Figure S3. Activation time and magnetic tweezers control experiments at 1 pN and 8.4 pN

(**A**) Distribution of the activation time, defined as the time until first unwinding event observation, in experiments using Flap-DNA at 1 pN in the presence of 100 nM HELB and 1 mM ATP. The distribution fits a Lorentzian function centred at 13 ± 6 min (n = 37).

(**B**) Representative time-courses of Flap-DNA control experiments performed in the absence of ATP at 1 pN force. (**C**) Representative time-courses of Flap-DNA control experiments performed in the presence of 100 nM HELB ATPase mutant at 1 pN. (**D**) Representative time-courses of experiments employing nicked DNA, done at 1 pN. (**E**) Distribution of the activation time in experiments using Flap-DNA at 8.4 pN in the presence of 100 nM HELB and 1 mM ATP. The distribution decays exponentially governed by a mean time <t> = 147 ± 10 s (n = 44). (**F**) Representative time-courses of Flap-DNA control experiments performed in the absence of ATP at 8.4 pN force. (**G**) Representative time-courses of Flap-DNA control experiments performed in the presence of 100 nM HELB ATPase mutant at 8.4 pN. (**H**) Representative time-courses of experiments employing nicked DNA, done at 8.4 pN. Shadowed regions indicate the time window of injection of protein's mixture.

**Figure S4. Dynamics of DNA loop formation by HELB at high force**

(**A**) Representative unwinding time-course obtained at 8.4 pN with Flap-DNA showing an unwinding (U) event and the formation and shrinkage of a loop. (**B**) Schematic representation of a model for DNA loop formation by HELB. Bead's height increases due to HELB unwinding of the duplex DNA by translocating from the 5′-ssDNA overhang. Movement of HELB is indicated by pink arrows. Exposure of a long ssDNA section will facilitate binding of additional HELB proteins, which will move with 5′-3′ polarity. A potential second binding site in HELB might facilitate the formation of a loop by keeping the protein still on the DNA. The formation of a loop results in a decrease of the bead's height. (**C**) The distribution of looping lengths decays exponentially with a mean length of 438 ± 60 bases (n = 40). (**D**) Distribution of the looping rate of events, including a Lorentzian function fit with mean velocity of 33 ± 14 base $s^{-1}$ (n = 37).

**Figure S5. HELB ssDNA-dependent activities are RPA species-specific**

(**A**) Bulk translocase assay based on displacement of streptavidin from 3′-biotinylated oligonucleotides. A time point was chosen (5 mins) where the reaction is not quite complete to be sensitive to changes induced by the presence of RPA. Increasing amounts of either human or yeast RPA were titrated to establish their effect on the translocation of HELB. (**B**) Quantification of the gel-based assay. HELB translocates at least as efficiently on ssDNA coated with human RPA. Under the same conditions, yeast RPA causes the activity to decrease significantly. (**C**) Change of ssDNA extension due to hRPA binding as a function of hRPA concentration at a constant force of 3.8 pN. (**D**) HELB translocation rate measured on hRPA-covered ssDNA at different hRPA concentrations. Experiments were performed at 5 nM HELB and 2 mM ATP. Box plots in **C** and **D** indicate the mean, median, 25$^{th}$ and 75$^{th}$ percentiles of the distributions and the whiskers show the standard deviation. Solid lines in **D** are normal distribution fits to the data. (**E**) Representative time-courses of MT experiments to study HELB activity on human RPA-coated ssDNA. Characteristic loop formation and shrinkage dynamics caused by HELB are observed. (**F**) Analogous MT experiments on yeast RPA coated ssDNA show no HELB activity. Shaded regions indicate the time window of reagents flow. Force is kept constant at 3.8 pN throughout the experiments. (**G**) RPA hinders HELB unwinding activity in bulk. HELB duplex-unwinding activity is fairly poor and appears to be inhibited by RPA, and to a greater extent than a generic bacterial SSB protein.

**Figure S6. HELB displacement of RPA is ATP- and species-specific**

(**A**) Representative kymograph showing movement of HELB-QD (blue) and removal of hRPA$^{MB543}$ (green) under ATP conditions (F = 15 pN). A schematic is overlaid showing the methodology to calculate HELB translocation rate ($\Delta L/\Delta t$). Two red arrows in merge panel point to two presumed non-labelled HELB trajectories. (**B**) (left panel) Representative merge kymograph showing HELB (blue) and ssDNA covered by hRPA$^{MB543}$ (green) in the absence of ATP. HELB binds but remains still in the absence of ATP (F = 19 pN). (right panel) Intensity profiles of the RPA$^{MB534}$ distribution at different times along the trajectory outlined by the coloured rectangles in the merge kymograph. Each line represents the average intensity over 10 frames (1.56 s). No change in the RPA distribution is detected. (**C**) Representative merge kymograph showing HELB (blue) and a ssDNA covered by yeast RPA$^{CY3}$ (green) in the presence of ATP. (**D**) Distribution of HELB translocation rates on yRPA-covered ssDNA. HELB translocation on ssDNA is mostly hindered by yeast RPA. Residual translocating events detected in long kymographs are fit to a Gaussian function with a peak at 67 ± 30 nt/s (n = 42).

**Supplementary Video 1. HELB translocation on ssDNA and RPA displacement** Video of a ssDNA tether between two optically-trapped beads covered by green fluorescent human RPA$^{MB543}$ (15 nM) showing the activity of biotinylated HELB labelled with QDs (5 nM HELB and 2 mM ATP, blue emission). The translocation of HELB towards the left bead is coupled to the displacement of RPA. F =  18 pN, pixel size = 50 nm, 5 fps.

#### SUPPLEMENTARY TABLES

##### Supplementary Table S1. Oligonucleotides used for fabrication of magnetic tweezers substrates.

| Fragment | Oligonucleotide | Sequence |
|---|---|---|
| Handles | 42.FMH_F2_KpnI-PsiI-ScaI | GCG TAA GTG GTA CCT TAT AAA GTA CTC GAC TCA CTA TAG GGA GAC CGG C |
| | JOE-R1 | AGT AAG CGC CGT CAG ACC AG |
| Flap | Poly(dT) flap | TTT TTT TTT TTT TTT TTT TTT TTT TTT TTT TTT TTT TTC AGC TAG CCT CAG CCT ACA ATC ACC |
| Complementary to released BbvCI fragment 1 | 137.block piece1 BbvCI | GGA TGA CAT GAG CTG A |
| Complementary to released BbvCI fragment2 | 138.block piece2 BbvCI | GGG TCA AGT GTG CTG A |
| Complementary to released BbvCI fragment3 | 139.block piece3 BbvCI | GGC TAG CTG AGC TGA |
| Complementary to released BbvCI fragment4 | 140.block piece4 BbvCI | GGT GAT TGT AGG CTG A |

##### Supplementary Table S2. Oligonucleotides used in translocase and helicase assays.

| Assay | Oligonucleotide | Sequence |
|---|---|---|
| Helicase assay | 3′ substrate | Cy5-GCT TGC TAG GAC GGA TCG CTC GAG GTT TTT TTT TTT TTT TTT TTT<br>+<br>C CTC GAG CGA TCC GTC CTA GCA AGC |
| | 5′ substrate | TTT TTT TTT TTT TTT TTT TTC CTC GAG CGA TCC GTC CTA GCA AGC<br>+<br>Cy5-GCT TGC TAG GAC GGA TCG CTC GAG G |
| Traslocation assay | 5′-3′ | 5′ - $^{32}$P G(biodT)ACGTATTCAAGATACCTCGTACTCTGTA CTGACTGATCCTAGG |
| | 3′-5′ | 5′ - $^{32}$P GTACGTATTCAAGATACCTCGTACTCTGTACTG ACTCGGATCC(biodT)A |
| EMSA assay | | Cy5-GCT TGC TAG GAC GGA TCG CTC GAG G |
| PIFE assay | | 3’-Cy3 labelled 15mer, 20mer, 30mer, 40mer poly(dT) |

**Supplementary Table S3. Sequence of DNA fragments used in this work**

Underlined sequence: 63 nucleotide gap created after digestion with the nicking enzyme Nt.BbvCI followed by denaturation. Sequence present in Flap-DNA substrate and Gap-DNA substrate.
Red underlined sequence: position of the Poly(dT) flap.
Black underlined sequence: position where the Poly(dT)-flap oligonucleotide anneals.

| Fragment | Size (bp) | Sequence |
|---|---|---|
| Central part of Flap-DNA substrate, Gap-DNA substrate and torsionally-constrained DNA substrates used in MT experiments. | 6337 | CAGTTCAGGAAGCGGTGATGCTGATAGAAGCCGGACTGAGTACCTACGAGAAA<br>GAGTGCGCAAAACGCGGTGACGACTATCAGGAAATTTTTGCCCAGCAGGTCCG<br>TGAAACGATGGAGCGCCGTGCAGCCGGTCTTAAACCGCCCGCCTGGGCGGCTG<br>CAGCATTTGAATCCGGGCTGCGACAATCAACAGAGGAGGAGAAGAGTGACAGC<br>AGAGCTGCGTAATCTCCCGCATATTGCCAGCATGGCCTTTAATGAGCCGCTGA<br>TGCTTGAACCCGCCTATGCGCGGGTTTTCTTTTGTGCGCTTGCAGGCCAGCTT<br>GGGATCAGCAGCCTGACGGATGCGGTGTCCGGCGACAGCCTGACTGCCCAGGA<br>GGCACTCGCGACGCTGGCATTATCCGGTGATGATGACGGACCACGACAGGCCC<br>GCAGTTATCAGGTCATGAACGGCATCGCCGTGCTGCCGGTGTCCGGCACGCTG<br>GTCAGCCGGACGCGGGCGCTGCAGCCGTACTCGGGGATGACCGGTTACAACGG<br>CATTATCGCCCGTCTGCAACAGGCTGCCAGCGATCCGATGGTGGACGGCATTC<br>TGCTCGATATGGACACGCCCGGCGGGATGGTGGCGGGGGCATTTGACTGCGCT<br>GACATCATCGCCCGTGTGCGTGACATAAAACCGGTATGGGCGCTTGCCAACGA<br>CATGAACTGCAGTGCAGGTCAGTTGCTTGCCAGTGCCGCCTCCCGGCGTCTGG<br>TCACGCAGACCGCCCGGACAGGCTCCATCGGCGTCATGATGGCTCACAGTAAT<br>TACGGTGCTGCGCTGGAGAAACAGGGTGTGGAAATCACGCTGATTTACAGCGG<br>CAGCCATAAGGTGGATGGCAACCCCTACAGCCATCTTCCGGATGACGTCCGGG<br>AGACACTGCAGTCCCGGATGGACGCAACCCGCCAGATGTTTGCGCAGAAGGTG<br>TCGGCATATACCGGCCTGTCCGTGCAGGTTGTGCTGGATACCGAGGCTGCAGT<br>GTACAGCGGTCAGGAGGCCATTGATGCCGGACTGGCTGATGAACTTGTTAACA<br>GCACCGATGCGATCACCGTCATGCGTGATGCACTGGATGCACGTAAATCCCGT<br>CTCTCAGGAGGGCGAATGACCAAAGAGACTCAATCAACAACTGTTTCAGCCAC<br>TGCTTCGCAGGCTGACGTTACTGACGTGGTGCCAGCGACGGAGGGCGAGAACG<br>CCAGCGCGGCGCAGCCGGACGTGAACGCGCAGATCACCGCAGCGGTTGCGGCA<br>GAAAACAGCCGCATTATGGGGATCCTCAACTGTGAGGAGGCTCACGGACGCGA<br>AGAACAGGCACGCGTGCTGGCAGAAACCCCCGGTATGACCGTGAAAACGGCCC<br>GCCGCATTCTGGCCGCAGCACCACAGAGTGCACAGGCGCGCAGTGACACTGCG<br>CTGGATCGTCTGATGCAGGGGGCACCGGCACCGCTGGCTGCAGGTAACCCGGC<br>ATCTGATGCCGTTAACGATTTGCTGAACACACCAGTGTAAGGGATGTTTATGA<br>CGAGCAAAGAAACCTTTACCCATTACCAGCCGCAGGGCAACAGTGACCCGGCT<br>CATACCGCAACCGCGCCCGGCGGATTGAGTGCGAAAGCGCCTGCAATGACCCC<br>GCTGATGCTGGACACCTCCAGCCGTAAGCTGGTTGCGTGGGATGGCACCACCG<br>ACGGTGCTGCCGTTGGCATTCTTGCGGTTGCTGCTCGAGCCTCAGCTCATGTC<br>ATCCTCAGCACACTTGACCCTCAGCTCAGCTAGCCTCAGCCTACAATCACCTC<br>AGCGAATTCGGTGACCCTTACGCGAATCCGCTTTCAGACGTTGACTGGTCGCG<br>TCTGGCAAAAGTTAAAGACCTGACGCCCGGCGAACTGACCGCTGAGTCCTATG<br>ACGACAGCTATCTCGATGATGAAGATGCAGACTGGACTGCGACCGGGCAGGGG<br>CAGAAATCTGCCGGAGATACCAGCTTCACGCTGGCGTGGATGCCCGGAGAGCA<br>GGGGCAGCAGGCGCTGCTGGCGTGGTTTAATGAAGGCGATACCCGTGCCTATA<br>AAATCCGCTTCCCGAACGGCACGGTCGATGTGTTCCGTGGCTGGGTCAGCAGT<br>ATCGGTAAGGCGGTGACGGCGAAGGAAGTGATCACCCGCACGGTGAAAGTCAC<br>CAATGTGGGACGTCCGTCGATGGCAGAAGATCGCAGCACGGTAACAGCGGCAA<br>CCGGCATGACCGTGACGCCTGCCAGCACCTCGGTGGTGAAAGGGCAGAGCACC<br>ACGCTGACCGTGGCCTTCCAGCCGGAGGGCGTAACCGACAAGAGCTTTCGTGC<br>GGTGTCTGCGGATAAAACAAAAGCCACCGTGTCGGTCAGTGGTATGACCATCA<br>CCGTGAACGGCGTTGCTGCAGGCAAGGTCAACATTCCGGTTGTATCCGGTAAT<br>GGTGAGTTTGCTGCGGTTGCAGAAATTACCGTCACCGCCAGTTAATCCGGAGA<br>GTCAGCGATGTTCCTGAAAACCGAATCATTTGAACATAACGGTGTGACCGTCA<br>CGCTTTCTGAACTGTCAGCCCTGCAGCGCATTGAGCATCTCGCCCTGATGAAA<br>CGGCAGGCAGAACAGGCGGAGTCAGACAGCAACCGGAAGTTTACTGTGGAAGA<br>CGCCATCAGAACCGGCGCGTTTCTGGTGGCGATGTCCCTGTGGCATAACCATC<br>CGCAGAAGACGCAGATGCCGTCCATGAATGAAGCCGTTAAACAGATTGAGCAG<br>GAAGTGCTTACCACCTGGCCCACGGAGGCAATTTCTCATGCTGAAAACGTGGT<br>GTACCGGCTGTCTGGTATGTATGAGTTTGTGGTGAATAATGCCCCTGAACAGA<br>CAGAGGACGCCGGGCCCGCAGAGCCTGTTTCTGCGGGAAAGTGTTCGACGGTG<br>AGCTGAGTTTTGCCCTGAAACTGGCGCGTGAGATGGGGCGACCCGACTGGCGT |

| | | |
|---|---|---|
| | | GCCATGCTTGCCGGGATGTCATCCACGGAGTATGCCGACTGGCACCGCTTTTA<br>CAGTACCCATTATTTTCATGATGTTCTGCTGGATATGCACTTTTCCGGGCTGA<br>CGTACACCGTGCTCAGCCTGTTTTTCAGCGATCCGGATATGCATCCGCTGGAT<br>TTCAGTCTGCTGAACCGGCGCGAGGCTGACGAAGAGCCTGAAGATGATGTGCT<br>GATGCAGAAAGCGGCAGGGCTTGCCGGAGGTGTCCGCTTTGGCCCGGACGGGA<br>ATGAAGTTATCCCCGCTTCCCCGGATGTGGCGGACATGACGGAGGATGACGTA<br>ATGCTGATGACAGTATCAGAAGGGATCGCAGGAGGAGTCCGGTATGGCTGAAC<br>CGGTAGGCGATCTGGTCGTTGATTTGAGTCTGGATGCGGCCAGATTTGACGAG<br>CAGATGGCCAGAGTCAGGCGTCATTTTTCTGGTACGGAAAGTGATGCGAAAAA<br>AACAGCGGCAGTCGTTGAACAGTCGCTGAGCCGACAGGCGCTGGCTGCACAGA<br>AAGCGGGGATTTCCGTCGGGCAGTATAAAGCCGCCATGCGTATGCTGCCTGCA<br>CAGTTCACCGACGTGGCCACGCAGCTTGCAGGCGGGCAAAGTCCGTGGCTGAT<br>CCTGCTGCAACAGGGGGGGCAGGTGAAGGACTCCTTCGGCGGGATGATCCCCA<br>TGTTCAGGGGGCTTGCCGGTGCGATCACCCTGCCGATGGTGGGGGCCACCTCG<br>CTGGCGGTGGCGACCGGTGCGCTGGCGTATGCCTGGTATCAGGGCAACTCAAC<br>CCTGTCCGATTTCAACAAAACGCTGGTCCTTTCCGGCAATCAGGCGGGACTGA<br>CGGCAGATCGTATGCTGGTCCTGTCCAGAGCCGGGCAGGCGGCAGGGCTGACG<br>TTTAACCAGACCAGCGAGTCACTCAGCGCACTGGTTAAGGCGGGGGTAAGCGG<br>TGAGGCTCAGATTGCGTCCATCAGCCAGAGTGTGGCGCGTTTCTCCTCTGCAT<br>CCGGCGTGGAGGTGGACAAGGTCGCTGAAGCCTCTAGAGAATGCTACGTACCT<br>GATGAGCTCCAGCTTTTGTTCCCTTTAGTGAGGGTTAATTGCGCGCTTGGCGT<br>AATCATGGTCATAGCTGTTTCCTGTGTGAAATTGTTATCCGCTCACAATTCCA<br>CACAACATACGAGCCGGAAGCATAAAGTGTAAAGCCTGGGGTGCCTAATGAGT<br>GAGCTAACTCACATTAATTGCGTTGCGCTCACTGCCCGCTTTCCAGTCGGGAA<br>ACCTGTCGTGCCAGCTGCATTAATGAATCGGCCAACGCGCGGGGAGAGGCGGT<br>TTGCGTATTGGGCGCTCTTCCGCTTCCTCGCTCACTGACTCGCTGCGCTCGGT<br>CGTTCGGCTGCGGCGAGCGGTATCAGCTCACTCAAAGGCGGTAATACGGTTAT<br>CCACAGAATCAGGGGATAACGCAGGAAAGAACATGTGAGCAAAAGGCCAGCAA<br>AAGGCCAGGAACCGTAAAAAGGCCGCGTTGCTGGCGTTTTTCCATAGGCTCCG<br>CCCCCCTGACGAGCATCACAAAAATCGACGCTCAAGTCAGAGGTGGCGAAACC<br>CGACAGGACTATAAAGATACCAGGCGTTTCCCCCTGGAAGCTCCCTCGTGCGC<br>TCTCCTGTTCCGACCCTGCCGCTTACCGGATACCTGTCCGCCTTTCTCCCTTC<br>GGGAAGCGTGGCGCTTTCTCATAGCTCACGCTGTAGGTATCTCAGTTCGGTGT<br>AGGTCGTTCGCTCCAAGCTGGGCTGTGTGCACGAACCCCCCGTTCAGCCCGAC<br>CGCTGCGCCTTATCCGGTAACTATCGTCTTGAGTCCAACCCGGTAAGACACGA<br>CTTATCGCCACTGGCAGCAGCCACTGGTAACAGGATTAGCAGAGCGAGGTATG<br>TAGGCGGTGCTACAGAGTTCTTGAAGTGGTGGCCTAACTACGGCTACACTAGA<br>AGGACAGTATTTGGTATCTGCGCTCTGCTGAAGCCAGTTACCTTCGGAAAAAG<br>AGTTGGTAGCTCTTGATCCGGCAAACAAACCACCGCTGGTAGCGGTGGTTTTT<br>TTGTTTGCAAGCAGCAGATTACGCGCAGAAAAAAAGGATCTCAAGAAGATCCT<br>TTGATCTTTTCTACGGGGTCTGACGCTCAGTGGAACGAAAACTCACGTTAAGG<br>GATTTTGGTCATGAGATTATCAAAAAGGATCTTCACCTAGATCCTTTTAAATT<br>AAAAATGAAGTTTTAAATCAATCTAAAGTATATATGAGTAAACTTGGTCTGAC<br>AGTTACCAATGCTTAATCAGTGAGGCACCTATCTCAGCGATCTGTCTATTTCG<br>TTCATCCATAGTTGCCTGACTCCCCGTCGTGTAGATAACTACGATACGGGAGG<br>GCTTACCATCTGGCCCCAGTGCTGCAATGATACCGCGAGACCCACGCTCACCG<br>GCTCCAGATTTATCAGCAATAAACCAGCCAGCCGGAAGGGCCGAGCGCAGAAG<br>TGGTCCTGCAACTTTATCCGCCTCCATCCAGTCTATTAATTGTTGCCGGGAAG<br>CTAGAGTAAGTAGTTCGCCAGTTAATAGTTTGCGCAACGTTGTTGCCATTGCT<br>ACAGGCATCGTGGTGTCACGCTCGTCGTTTGGTATGGCTTCATTCAGCTCCGG<br>TTCCCAACGATCAAGGCGAGTTACATGATCCCCCATGTTGTGCAAAAAAGCGG<br>TTAGCTCCTTCGGTCCTCCGATCGTTGTCAGAAGTAAGTTGGCCGCAGTGTTA<br>TCACTCATGGTTATGGCAGCACTGCATAATTCTCTTACTGTCATGCCATCCGT<br>AAGATGCTTTTCTGTGACTGGTGAGTACTCAACCAAGTCATTCTGAGAATAGT<br>GTATGCGGCGACCGAGTTGCTCTTGCCCGGCGTCAATACGGGATAATACCGCG<br>CCACATAGCAGAACTTTAAAAGTGCTCATCATTGGAAAACGTTCTTCGGGGCG<br>AAAACTCTCAAGGATCTTACCGCTGTTGAGATCCAGTTCGATGTAACCCACTC<br>GTGCACCCAACTGATCTTCAGCATCTTTTACTTTCACCAGCGTTTCTGGGTGA<br>GCAAAAACAGGAAGGCAAAATGCCGCAAAAAAGGGAATAAGGGCGACACGGAA<br>ATGTTGAATACTCATACTCTTCCTTTTTCAATATTATTGAAGCATTTATCAGG<br>GTTATTGTCTCATGAGCGGATACATATTTGAATGTATTTAGAAAAATAAACAA<br>ATAGGGGTTCCGCGCACATTTCCCCGAAAAGTGCCACCTAAATTGTAAGCGTT<br>AATATTTTGTTAAAATTCGCGTTAAATTTTTGTTAAATCAGCTCATTTTTTAA<br>CCAATAGGCCGAAATCGGCAAAATCCCTTA |

**Figure S1**

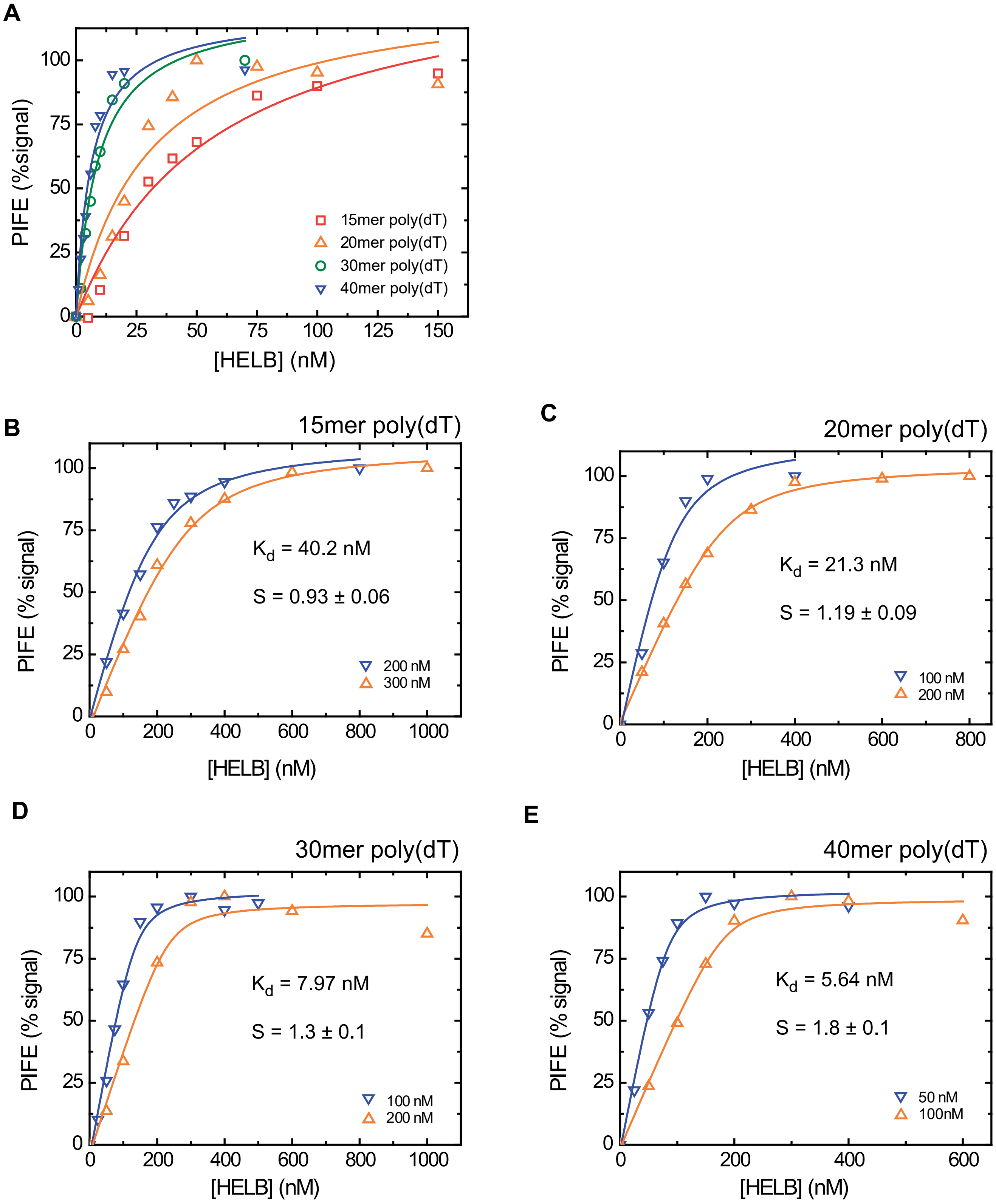

A
PIFE (%signal)
[HELB] (nM)
15mer poly(dT)
20mer poly(dT)
30mer poly(dT)
40mer poly(dT)
B
15mer poly(dT)
PIFE (% signal)
[HELB] (nM)
K$_d$ = 40.2 nM
S = 0.93 ± 0.06
200 nM
300 nM
C
20mer poly(dT)
PIFE (% signal)
[HELB] (nM)
K$_d$ = 21.3 nM
S = 1.19 ± 0.09
100 nM
200 nM
D
30mer poly(dT)
PIFE (% signal)
[HELB] (nM)
K$_d$ = 7.97 nM
S = 1.3 ± 0.1
100 nM
200 nM
E
40mer poly(dT)
PIFE (% signal)
[HELB] (nM)
K$_d$ = 5.64 nM
S = 1.8 ± 0.1
50 nM
100nM

**Figure S2**

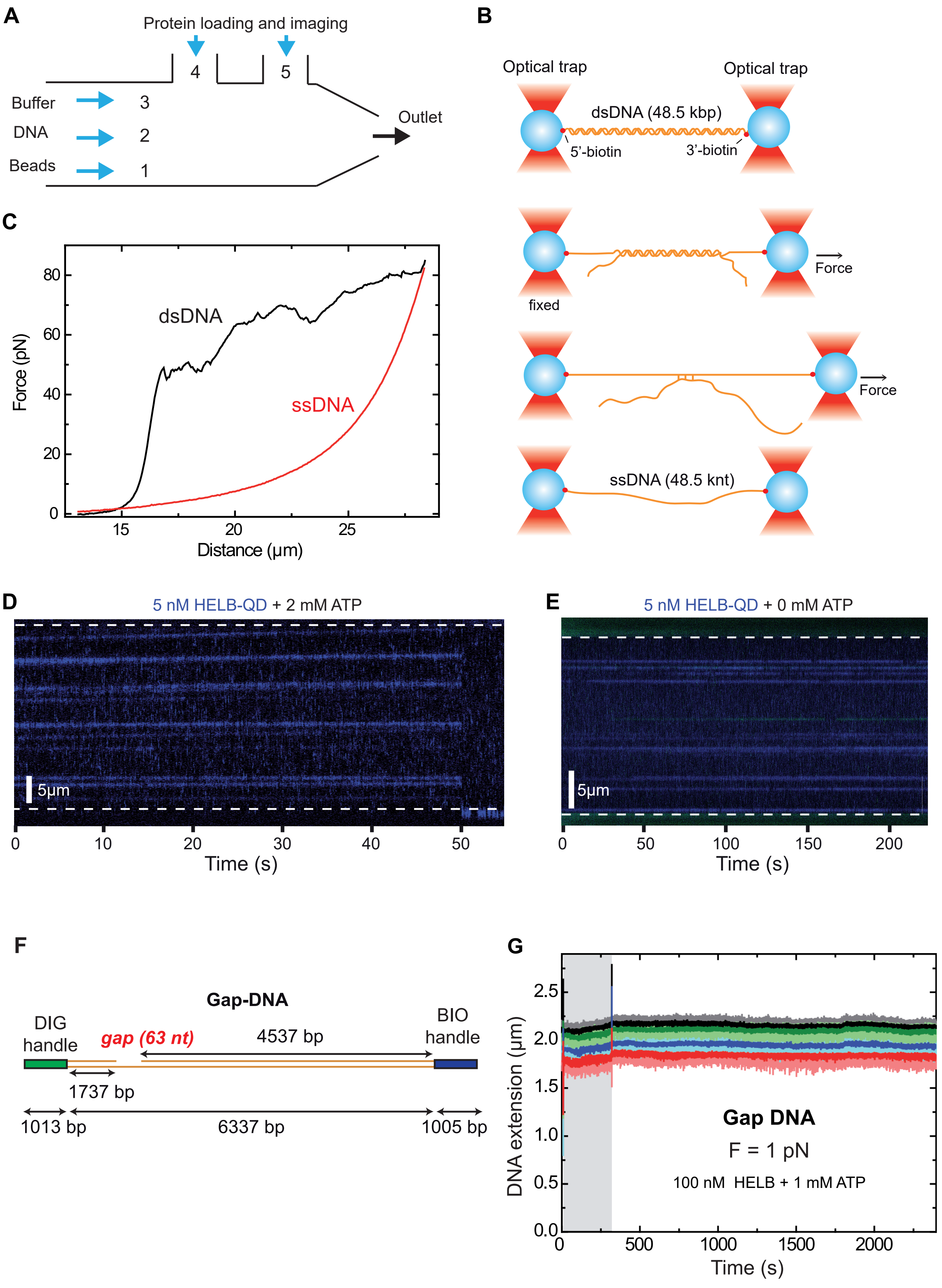
A
Protein loading and imaging
4
5
Buffer
3
DNA
2
Beads
1
Outlet
B
Optical trap
Optical trap
dsDNA (48.5 kbp)
5'-biotin
3'-biotin
Force
fixed
Force
ssDNA (48.5 knt)
C
dsDNA
ssDNA
Force (pN)
Distance (µm)
D
5 nM HELB-QD + 2 mM ATP
5µm
Time (s)
E
5 nM HELB-QD + 0 mM ATP
5µm
Time (s)
F
Gap-DNA
DIG
handle
gap (63 nt)
4537 bp
BIO
handle
1737 bp
1013 bp
6337 bp
1005 bp
G
DNA extension (µm)
Gap DNA
F = 1 pN
100 nM HELB + 1 mM ATP
Time (s)

**Figure S3**

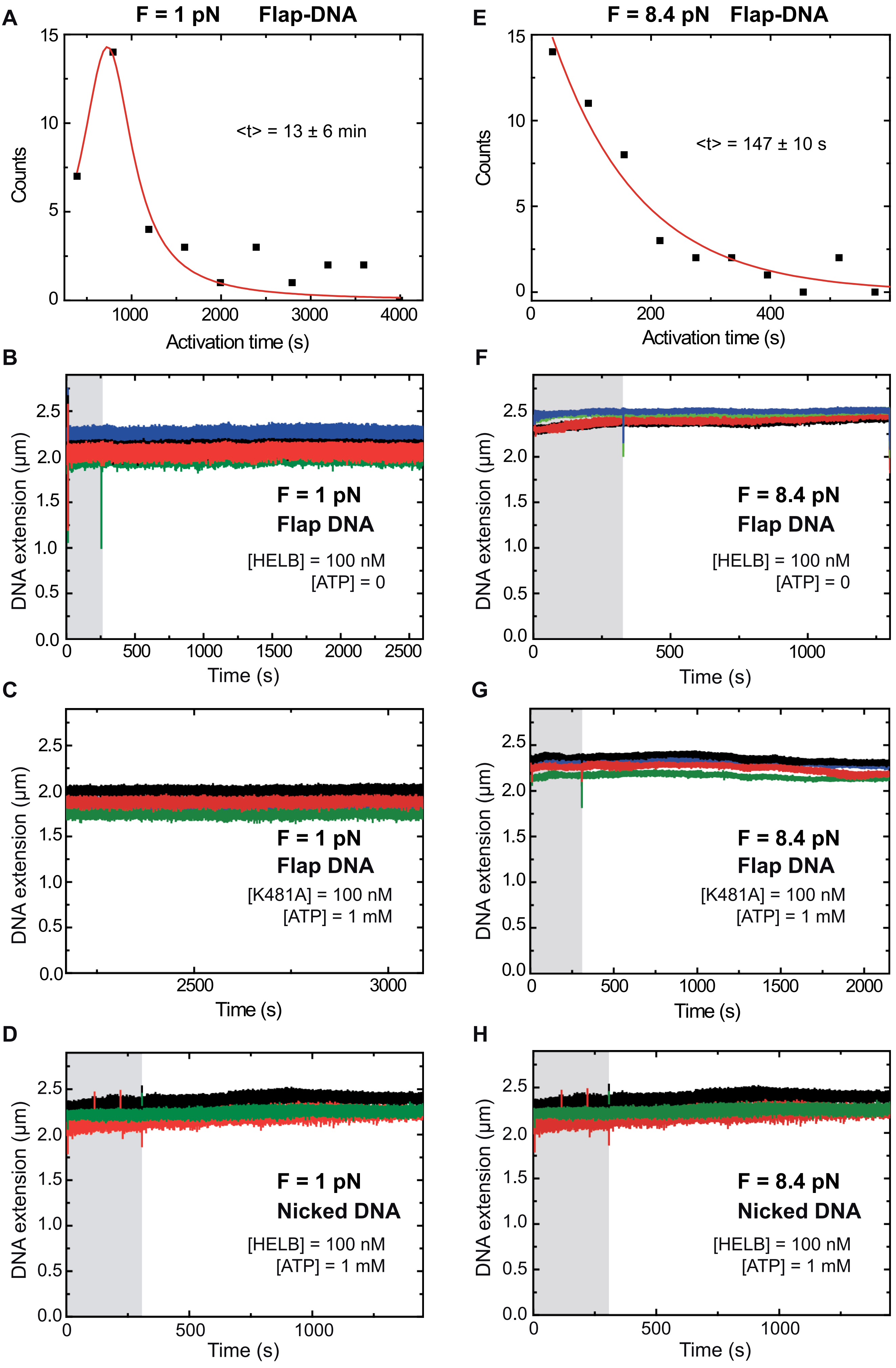
A
F = 1 pN Flap-DNA
<t> = 13 ± 6 min
Counts
Activation time (s)
E
F = 8.4 pN Flap-DNA
<t> = 147 ± 10 s
Counts
Activation time (s)
B
F = 1 pN
Flap DNA
[HELB] = 100 nM
[ATP] = 0
DNA extension (µm)
Time (s)
F
F = 8.4 pN
Flap DNA
[HELB] = 100 nM
[ATP] = 0
DNA extension (µm)
Time (s)
C
F = 1 pN
Flap DNA
[K481A] = 100 nM
[ATP] = 1 mM
DNA extension (µm)
Time (s)
G
F = 8.4 pN
Flap DNA
[K481A] = 100 nM
[ATP] = 1 mM
DNA extension (µm)
Time (s)
D
F = 1 pN
Nicked DNA
[HELB] = 100 nM
[ATP] = 1 mM
DNA extension (µm)
Time (s)
H
F = 8.4 pN
Nicked DNA
[HELB] = 100 nM
[ATP] = 1 mM
DNA extension (µm)
Time (s)

Figure S4

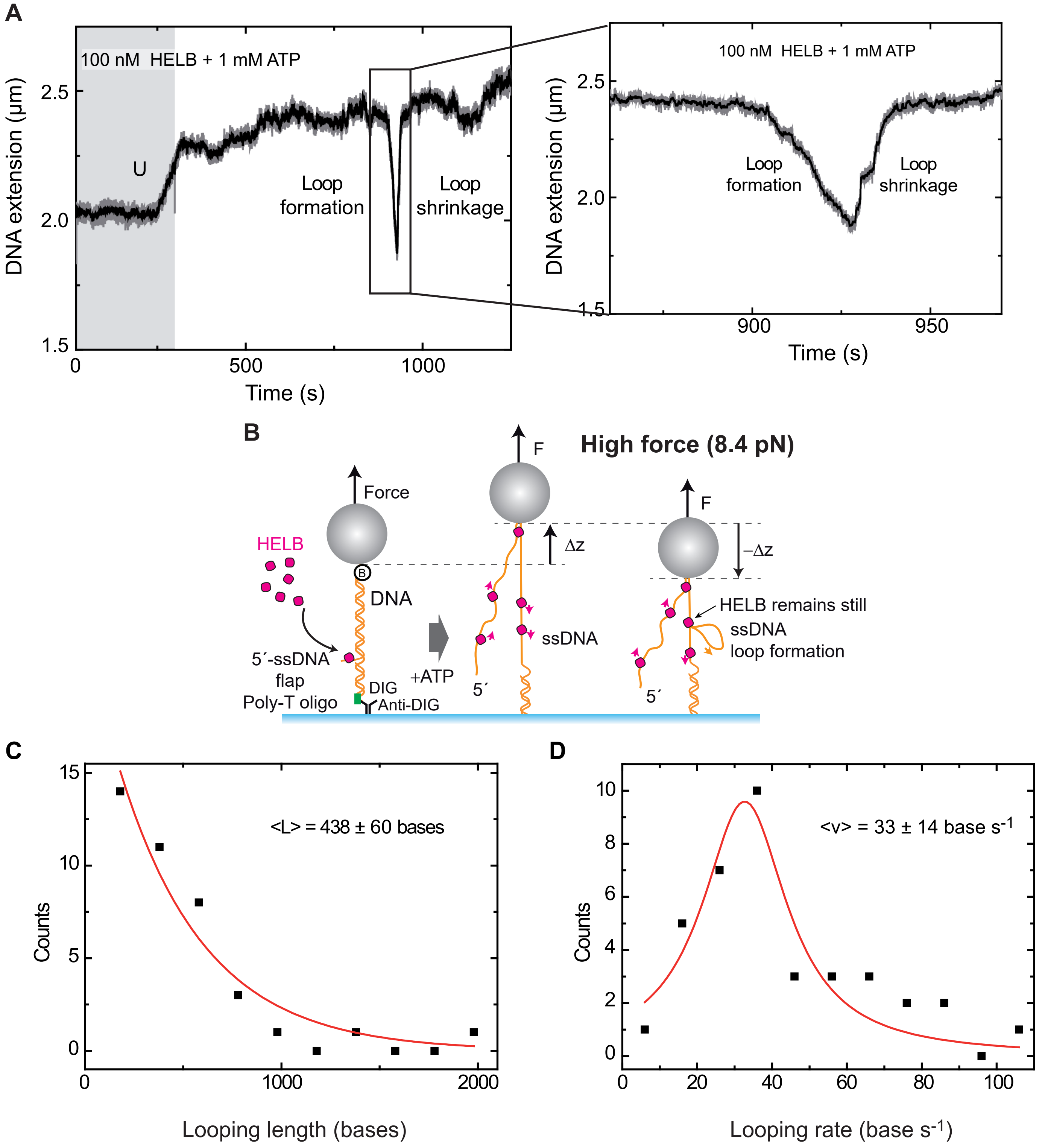
A
100 nM HELB + 1 mM ATP
DNA extension (µm)
U
Loop formation
Loop shrinkage
Time (s)
B
High force (8.4 pN)
Force
F
HELB
DNA
Δz
−Δz
HELB remains still
ssDNA
ssDNA loop formation
5´-ssDNA flap
Poly-T oligo
DIG
Anti-DIG
+ATP
5´
C
Counts
<L> = 438 ± 60 bases
Looping length (bases)
D
Counts
<v> = 33 ± 14 base s-1
Looping rate (base s-1)

### Figure S5

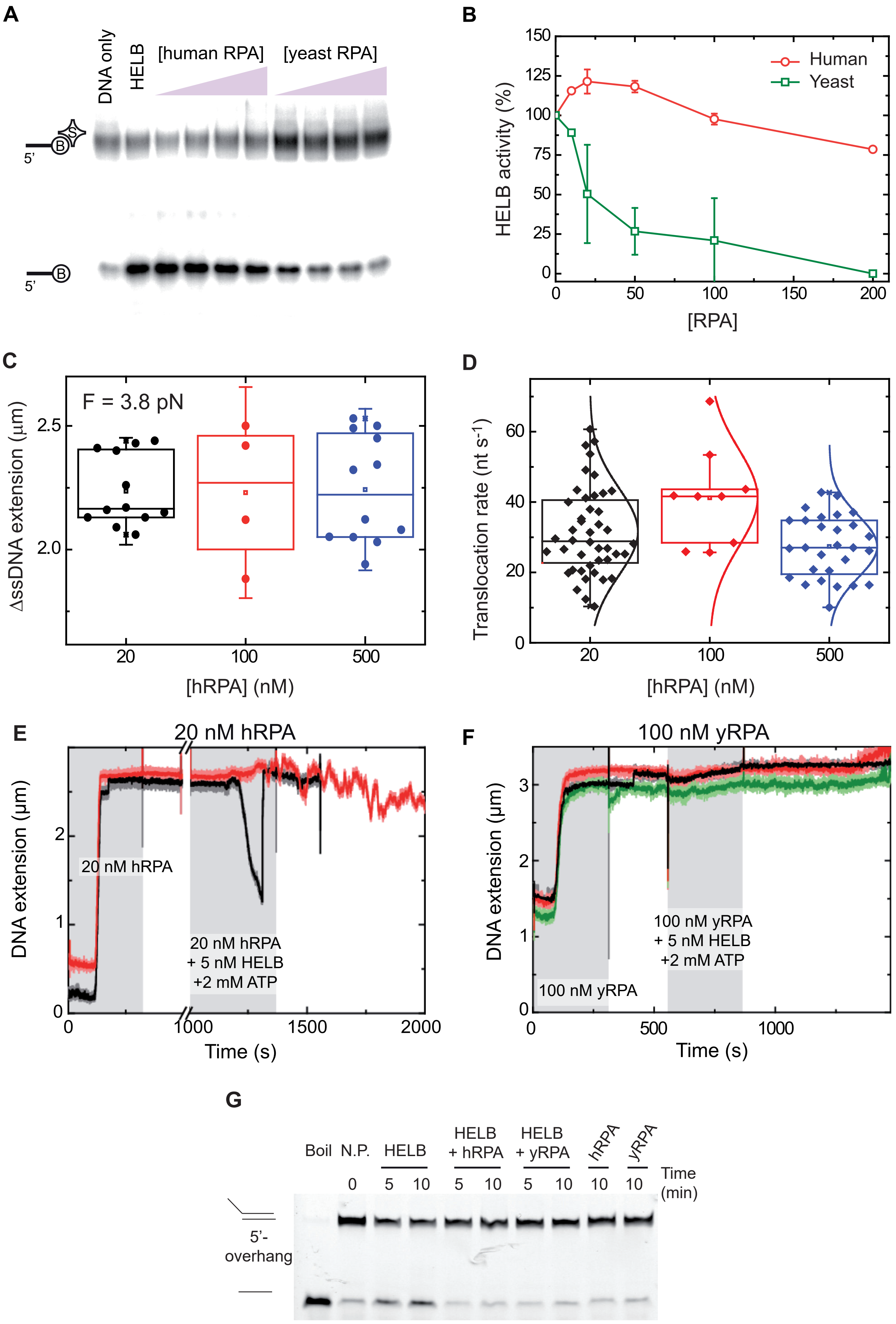
A
DNA only
HELB
[human RPA]
[yeast RPA]
5'
5'
B
Human
Yeast
HELB activity (%)
[RPA]
C
F = 3.8 pN
ΔssDNA extension (μm)
[hRPA] (nM)
D
Translocation rate (nt s-1)
[hRPA] (nM)
E
20 nM hRPA
20 nM hRPA
20 nM hRPA + 5 nM HELB +2 mM ATP
DNA extension (μm)
Time (s)
F
100 nM yRPA
100 nM yRPA
100 nM yRPA + 5 nM HELB +2 mM ATP
DNA extension (μm)
Time (s)
G
Boil
N.P.
HELB
HELB + hRPA
HELB + yRPA
hRPA
yRPA
Time (min)
5'-overhang

**Figure S6**

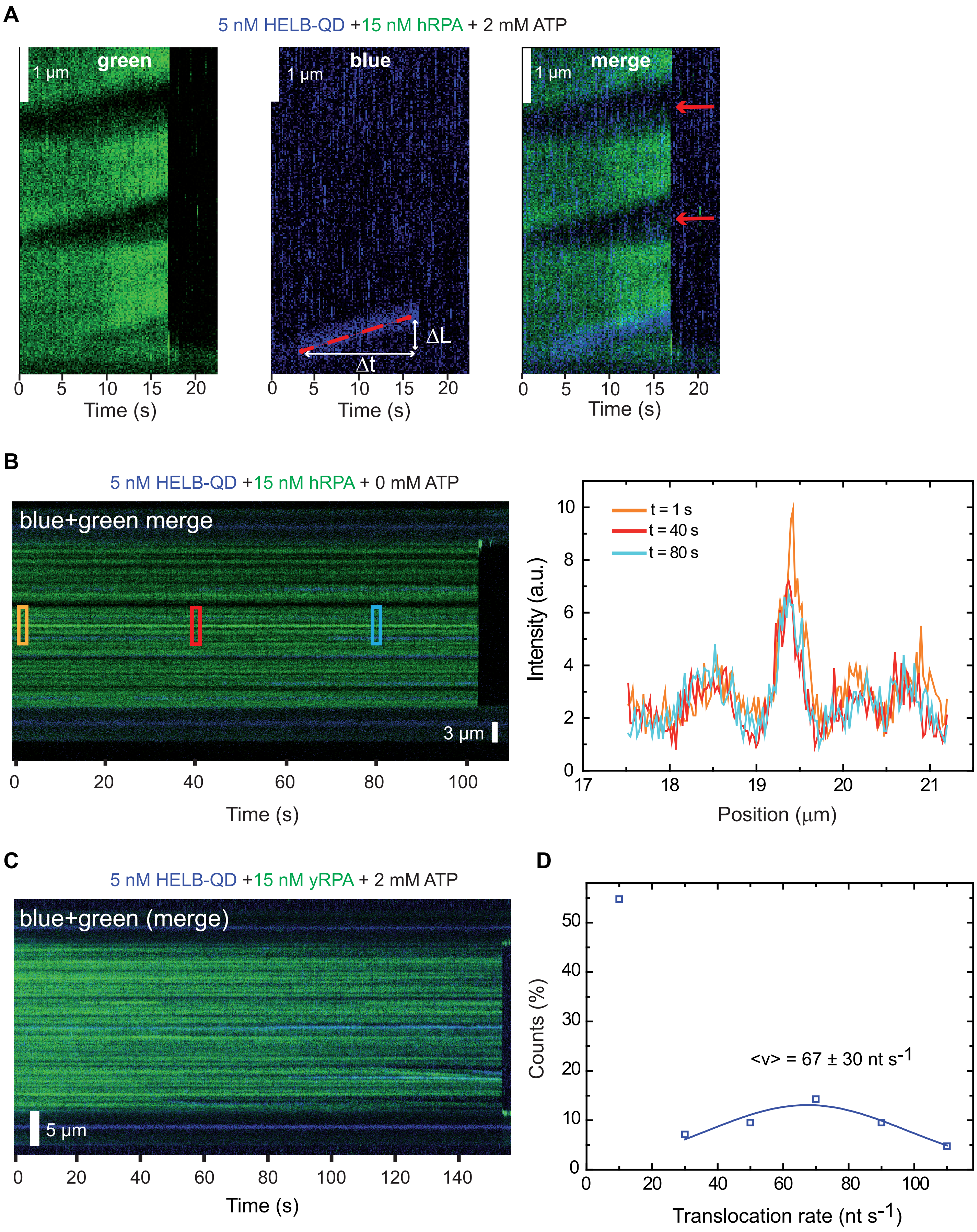
A
5 nM HELB-QD +15 nM hRPA + 2 mM ATP
green
1 µm
blue
1 µm
merge
1 µm
ΔL
Δt
0 5 10 15 20
Time (s)
B
5 nM HELB-QD +15 nM hRPA + 0 mM ATP
blue+green merge
3 µm
0 20 40 60 80 100
Time (s)
t = 1 s
t = 40 s
t = 80 s
Intensity (a.u.)
Position (µm)
C
5 nM HELB-QD +15 nM yRPA + 2 mM ATP
blue+green (merge)
5 µm
0 20 40 60 80 100 120 140
Time (s)
D
Counts (%)
<v> = 67 ± 30 nt s-1
Translocation rate (nt s-1)
